# Cell autonomous requirement of Neurofibromin (Nf1) for postnatal muscle hypertrophic growth and metabolic homeostasis

**DOI:** 10.1101/2020.04.24.059931

**Authors:** Xiaoyan Wei, Julia Franke, Mario Ost, Kristina Wardelmann, Stefan Börno, Bernd Timmermann, David Meierhofer, Andre Kleinridders, Susanne Klaus, Sigmar Stricker

**Affiliations:** Musculoskeletal Development and Regeneration Group, Institute of Chemistry and Biochemistry, Freie Universität Berlin, Berlin, Germany; Development and Disease Group, Max Planck Institute for Molecular Genetics, Berlin, Germany; Department of Physiology of Energy Metabolism, German Institute for Human Nutrition, Nuthetal, Germany; Department of Neuropathology, University Hospital Leipzig, Leipzig, Germany; Junior Research Group Central Regulation of Metabolism, German Institute for Human Nutrition, Nuthetal, Germany; German Center for Diabetes Research (DZD), D-85764 München-Neuherberg, Germany; Sequencing Core Unit, Max Planck Institute for Molecular Genetics, Berlin, Germany; Mass Spectrometry Core Unit, Max Planck Institute for Molecular Genetics, Berlin, Germany; University of Potsdam, Institute of Nutritional Science, Potsdam, Germany

**Keywords:** Neurofibromatosis, myopathy, muscle atrophy, muscle metabolism, muscle fiber type, AMPK

## Abstract

**Background:** Neurofibromatosis type 1 (NF1) is a multi-organ disease caused by mutations in Neurofibromin *(NF1).* Amongst other features, NF1 patients frequently show reduced muscle mass and strength, impairing patients’ mobility and increasing the risk of fall. The role of Nf1 in muscle and the cause for the NF1-associated myopathy is mostly unknown.

**Methods:** To dissect the function of Nf1 in muscle, we created muscle-specific knockout mouse models for Nf1, inactivating Nf1 in the prenatal myogenic lineage either under the Lbx1 promoter or under the Myf5 promoter. Mice were analyzed during pre-and postnatal myogenesis and muscle growth.

**Results:** Nf1^Lbx1^ and Nf1^Myf5^ animals showed only mild defects in prenatal myogenesis. Nf1^Lbx1^ animals were perinatally lethal, while Nf1^Myf5^ animals survived only up to approx. 25 weeks. A comprehensive phenotypic characterization of Nf1^Myf5^ animals showed decreased postnatal growth, reduced muscle size, and fast fiber atrophy. Proteome and transcriptome analysis of muscle tissue indicated decreased protein synthesis and increased proteasomal degradation, and decreased glycolytic and increased oxidative activity in muscle tissue. High-resolution respirometry confirmed enhanced oxidative metabolism in Nf1^Myf5^ muscles, which was concomitant to a fiber type shift from type 2B to type 2A and type 1. Moreover, Nf1^Myf5^ muscles showed hallmarks of decreased activation of mTORC1 and increased expression of atrogenes. Remarkably, loss of Nf1 promoted a robust activation of AMPK with a gene expression profile indicative of increased fatty acid catabolism. Additionally, we observed a strong induction of genes encoding catabolic cytokines in muscle Nf1^Myf5^ animals, in line with a drastic reduction of white, but not brown adipose tissue.

**Conclusions:** Our results demonstrate a cell-autonomous role for Nf1 in myogenic cells during postnatal muscle growth required for metabolic and proteostatic homeostasis. Furthermore, Nf1 deficiency in muscle drives cross-tissue communication and mobilization of lipid reserves.

## Introduction

Neurofibromatosis type I (NF1) is a multi-organ disease caused by mutations in the *NF1* gene. *NF1* encodes a Ras-GTPase-activating protein (Ras-GAP), Neurofibromin, which negatively regulates Ras signaling, mainly feeding into the MAPK pathway, specifically Mek1/2 – Erk1/2 signaling, depending on the cell type. In contrast, other pathways were shown to be activated [1]. The musculoskeletal system is often affected in NF1 patients, with scoliosis, long bone dysplasia and osteoporosis being a main cause of considerable morbidity [2]. Furthermore, pronounced reduction of muscle size and muscle weakness has been reported in NF1 patients [3–8], and has also been described as general feature in RASopathies, i.e. disease traits with mutations in RAS pathway components, [9,10]. However, the cause of muscle weakness is unknown.

Skeletal muscles consist of syncytial multinucleated myofibers that produce force based on the activity of contractile proteins. During postnatal growth, myofibers continuously increase their size by protein synthesis, mainly components of the contractile machinery, in a process termed hypertrophy. Myofibers are generally classified into slow twitching type 1 and fast twitching type 2 fibers that differ in the expression of specific myosin heavy chain isoforms as well as their metabolic profile. While slow twitching fibers mainly rely on oxidative metabolism based on carbohydrates and fatty acids, fast twitch fibers are further divided into fast oxidative (type 2A) or fast glycolytic (type 2B and 2X in mice, 2X in humans) fibers [11–13]. Myopathies affect the functionality of muscle fibers, by e.g. perturbing the function of contractile proteins or the integrity of the myofiber, or by changes in protein turnover leading to net protein loss and myofiber atrophy. Thereby, fiber types can be differentially affected, e.g. in Duchenne muscular dystrophy or in age-related sarcopenia predominantly fast glycolytic fibers are most prone to atrophy [13,14]

During development, skeletal muscles of the trunk arise from the somitic mesoderm. The muscles of the limbs instead originate from the ventrolateral lip of the dermomyotome, where cells detach and migrate into the limb buds, where they invade the resident mesenchyme. There, cells start to induce a cascade of so-called myogenic regulatory factors (Myf5, Myf6/Mrf4, Myod, Myogenin) that consecutively specify and consolidate myogenic commitment and differentiation. Inactivation of *Nf1* in limb mesenchyme during development using Prx1^Cre^ recapitulated the skeletal manifestations of NF1 [15,16], but also caused an early-onset defect in embryonal myogenesis [17], providing the first evidence for a direct involvement of Nf1 in myogenesis. Postnatally, Nf1^Prx1^ mice recapitulated Nf1 features as muscle weakness and fibro-fatty infiltration of muscle [17,18]. However, in this model, Nf1 is inactivated in the skeleton as well as in the accessory tissues of muscles (tendons, connective tissue) [17,19]. Inactivation of Nf1 in the myogenic lineage using Myod^Cre^ resulted in early postnatal lethality [20] limiting the utility of this model for postnatal analysis.

Here we use Lbx1^Cre^ and Myf5^Cre^ to delete Nf1 in the myogenic lineage. Lbx1^Cre^ leads to inactivation in early migrating progenitors, while Myf5^Cre^ leads to inactivation in committed myoblasts. While Lbx1^Cre^-mediated inactivation of Nf1 led to perinatal lethality, inactivation of Nf1 via Myf5^Cre^ provided viable offspring and allowed postnatal analysis. We show that loss of Nf1 in the myogenic lineage compromises postnatal muscle growth resulting in severely reduced muscle size caused by decreased mTORC1 signaling activity. Moreover, Nf1 deficient muscle shows a glycolytic-to-oxidative fiber type shift concomitant to decreased carbohydrate, but increased lipid metabolism and attrition of white adipose tissue. This demonstrates a cell-autonomous requirement for Nf1 in the myogenic lineage and provides a mechanistic explanation for Nf1-associated muscle weakness.

## Methods

### Animals

Mouse lines used in this study were described before; Myf5^Cre^ [21] Lbx1^Cre^ [22] Nf1^flox^ [23] and Rosa26^mTmG^ [24]. All animal procedures conducted within this study were conducted in accordance with FELASA guidelines and were approved by the responsible authority (Landesamt für Gesundheit und Soziales Berlin, LaGeSo) under license numbers ZH120, G0346/13, and G0176/19. Timed matings were set up, and mice were sacrificed by cervical dislocation at indicated days of pregnancy, fetuses were sacrificed by decapitation.

### Tissue Processing and Histology

Forelimbs and hind limbs of mice were dissected, limbs from the right side of the body were embedded in paraffin and limbs from the other side snap-frozen in liquid nitrogen, followed by RNA, gDNA or protein isolation. Adipose tissue was taken for RNA isolation and paraffin embedding. Muscle tissue was embedded with tragacanth (Sigma-Aldrich #G1128) on a cork plate into cold isopentane for 10 seconds then samples were transferred on dry ice and stored at −80°C. Adipose tissue was embedded with paraffin after using PFA fixation and rounds of dehydration. Cryoembedded samples were sectioned using a Microtome (Microm HM355S) with a thickness of 10 μm. For immunostaining slides were fixed with methanol for 10 min. at −20 °C. Antigen retrieval was used for Pax7 staining with antigen retrieval solution (2mM EDTA) for 10min at 95°C water bath (Julabo). Slides were blocked with blocking buffer (5% BSA in PBX) for 1h at room temperature. Primary antibodies were diluted with blocking buffer and incubated overnight at 4 °C. To remove primary antibody slides were washed 4 times 10 min each with PBX. Secondary antibody was applied in PBX for 1h at room temperature followed by washing with PBX for 4 times. Slides were mounted with Fluoromount-G (Southern Biotech, #0100-01). Imaging was performed on a LSM700 confocal microscope (Zeiss) with ZEN imaging software (Zeiss). The primary antibodies and dilutions used were: Goat anti-Collagen IV (Millipore AB769; 1:500), Mouse anti-MyHC Type 1 (DSHB BA-D5; 1:500), Mouse anti-MyHC Type2A (DSHB SC-71; 1:200), Rabbit anti-Laminin (Sigma-Aldrich L9393; 1:500), Mouse anti-Pax7 (DSHB Pax7; 1:10), Mouse anti-Myosin Heavy Chain (Sigma-Aldrich 05-716; 1:1000), Anti-beta III tubulin antibody (Sigma-Aldrich AB9354; 1:500).

For Hematoxylin & Eosin staining, slides were fixed with 4 % PFA at room temperature for 10 min, then incubated with hematoxylin (Carl Roth 3816) for 1 min, followed by eosin (Sigma Aldrich # 1508694-9) staining for 30 sec. Slides were dehydrated by incubation for 5 min each in 70% EtOH–80% EtOH – 90% EtOH – 100% EtOH – Xylene (Carl Roth #CN80). Slides were mounted with permount medium (Science Service E17986).

For Oil Red O staining, slides were rinsed with 60% isopropanol (Carl Roth 6752) for 10 min followed by incubation with Oil Red O (Sigma Aldrich O0625) staining solution (0.5 g Oil Red O in 100 ml isopropanol) for 15 min. Then slides were rinsed with 60% isopropanol for 10 min and washed with PBS before mounting and imaging.

### RT-q-PCR

Total RNA was isolated with RNeasy Micro Kit or RNeasy Mini Kit (Qiagen). cDNA was synthesized using 1μg of mRNA with SuperScript™ III Reverse Transcriptase (Invitrogen 18080044), Oligo(dT)20 Primer (Invitrogen 18418020) and RNaseOUT Recombinant Ribonuclease Inhibitor (ThermoFisher Scientific 10777019) according to the protocol provided by the manufactures. In brief, 1:10 cDNA dilution was used for each reaction of quantitative RT-PCR analysis performed on an ABI Prism HT 7900 real-time PCR detection system (Applied Biosystems) equipped with SDS software version 2.4 (ThermoFisher Scientific) using GOTaq qPCR Master Mix (Promega) or SYBR Green qPCR Master Mix (Life Technologies). Beta-actin was used to normalize the expression of each gene. All primers used were purchased from Eurofins Scientific and are listed in the supporting information, Table S1.

### RNA sequencing analysis

Total RNA was isolated using RNeasy Micro Kit or RNeasy Mini Kit (Qiagen). After initial quality control using Agilent’s Bioanalyzer sequencing libraries were prepared from 500ng of total RNA per sample following Roche’s “KAPA stranded mRNA Seq” library preparation protocol for single indexed Illumina libraries: First the polyA-RNA fraction was enriched using oligo-dT-probed paramagnetic beads. Enriched RNA was heat-fragmented and subjected to first strand synthesis using random priming. The second strand was synthesized incorporating dUTP instead of dTTP to preserve strand information. After A-tailing Illumina sequencing compatible adapters were ligated. Following bead-based clean-up steps the libraries were amplified using 11 cycles of PCR. Library quality and size was checked with qBit, Agilent Bioanalyzer and qPCR. Sequencing was carried out on an Illumina HiSeq 4000 system in PE75bp mode yielding between 45-72 million fragments per sample. Mapping was performed using STAR 2.4.2a software with mouse genome (mm9). Read counts were generated with R Studio function Summarize Overlaps and normalized to RPKM based on the number of uniquely mapped reads. Differential expression analysis was performed with DESeq2 using default settings. Genes with an absolute fold change>2 and adjusted p-value<0.01 were assessed to be significantly differentially expressed. A table of DESeq2 analysis can be found in the supporting information, Data S1. GSEA analysis was performed with the entire gene list using GSEA software 4.0.1 desktop (Broad Institute). KEGG and Gene ontology enrichment analysis was performed using the Signature Database (MSigDB) and DAVID 6.8. RNA-seq data have been deposited in the Gene Expression Omnibus (GEO) database under the SuperSeries accession number GSE147605.

### Immunoblotting

For protein isolation, tissue homogenization was performed with a TissueLyser (Qiagen) with RIPA buffer (50 mM Tris-HCl, pH 8.0; 150 mM NaCl; 1% NP-40; 0.5% Sodium deoxycholate; 0.1% SDS). Protein concentration was measured with the Pierce BCA Protein Assay Kit (Thermo Fischer #23225). Total protein was separated with SDS-PAGE gels and transferred to PVDF membrane (GE Healthcare). Membranes were blocked with 5% BSA in TBST for 1h at room temperature. Primary antibodies were diluted in blocking buffer and applied overnight at 4° C. Antibodies used: Rabbit anti-phosphor (Thr 389) 70s6k (Cell Signaling Technology 9205), Rabbit anti-Pparg (Cell Signaling Technology 2443), Rabbit anti-phosphor (pThr172) AMPKa (Cell Signaling Technology 2535), Rabbit anti-AMPKα (Cell Signaling Technology 2532), Mouse anti-β-actin (Cell Signaling Technology 58169), Mouse anti-β-tubulin (Sigma-Aldrich T8328), Rabbit anti-phosphor (s235/236) S6 (Cell Signaling Technology 4858), Mouse anti-MyHC Type 1 (DSHB BA-D5), Mouse anti-MyHc Type2B (DSHB BF-F3), Rabbit anti-phosphor (ser 473) AKT (Cell signaling, # 9271; 1:1000), Rabbit anti-Phospho (Ser1101) IRS-1(Cell signaling, #2385; 1:1000), Rabbit anti-Phospho (Ser636/639) IRS-1(Cell signaling, # 2388; 1:1000). Secondary antibodies were applied for 1 hour at room temperature. Antibodies used: IgG (HRP Conjugated) anti-Rabbit (Thermo Fisher, 1:1000), IgG (HRP Conjugated) anti-Mouse (Thermo Fisher, 1:1000). Images were acquired using a Fusion FX spectra gel documentation system (Vilber) with FUSION FX software. Blots images were analyzed with ImageJ gray value analysis tool.

### Proteomics sample preparation

Three biological replicates of Nf1^Myf5^ and control (Nf1^flox/+^;Myf5^Cre^ and Nf1^+/+^;Myf5^Cre^) TA muscles from p21 mice were used for proteome profiling. Proteomics sample preparation was done according to a published protocol with minor modifications [25]. In brief, about 10 mg frozen tissue per sample were homogenized under denaturing conditions with a FastPrep (2 times for 60 s, 4 m x s-1) in 1 ml of a buffer containing 6 M guanidinium chloride (GdmCl), 5 mM tris(2-carboxyethyl)phosphine, 20 mM chloroacetamide and 50 mM Tris-HCl pH 8.5. Lysates were boiled at 95°C for 15 min in a thermal shaker, followed by sonication for 15 min and centrifuged at 15,000 g for 5 min. The supernatant was transferred into new protein low binding tubes (Eppendorf, Germany). 100 μg protein per sample were diluted to 1 M GdmCl by adding 10% acetonitrile and 25 mM Tris, 8.5 pH, followed by a Lys C digestion (1:50) at 37°C for 2 hours. This was followed by another dilution to 0.5 M GdmCl and a tryptic digestion (1:50) at 37°C overnight. Subsequently, peptides were desalted with C18 columns and reconstituted in 1% formic acid in water and further separated into five fractions by strong cation exchange chromatography (SCX, 3M Purification, Meriden, CT). Eluates were first dried in a SpeedVac, then dissolved in 20 μl 5% acetonitrile and 2% formic acid in water, briefly vortexed, and sonicated in a water bath for 30 seconds prior injection to nano-LC-MS/MS. Five μg of each SCX fraction and a non-fractioned sample were used for proteome profiling and analyzed by MaxQuant (v1.5.3.30).

### LC-MS/MS instrument settings for shotgun proteome profiling with Label-FreeQuantification (LFQ) and data analysis

LC-MS/MS was carried out by nanoflow reverse phase liquid chromatography (Dionex Ultimate 3000, Thermo Scientific) coupled online to a Q-Exactive HF Orbitrap mass spectrometer (Thermo Scientific), as reported previously [26]. Briefly, the LC separation was performed using a PicoFrit analytical column (75 μm ID × 50 cm long, 15 μm Tip ID; New Objectives, Woburn, MA) in-house packed with 3-μm C18 resin (Reprosil-AQ Pur, Dr. Maisch, Ammerbuch, Germany). Peptides were eluted using a gradient from 3.8 to 38% solvent B in solvent A over 120 min at 266 nL per minute flow rate. Solvent A was 0.1 % formic acid and solvent B was 79.9% acetonitrile, 20% H_2_O, 0.1% formic acid. Nanoelectrospray was generated by applying 3.5 kV. A cycle of one full Fourier transformation scan mass spectrum (300-1750 m/z, resolution of 60,000 at m/z 200, automatic gain control (AGC) target 1 × 10^6^) was followed by 12 data-dependent MS/MS scans (resolution of 30,000, AGC target 5 × 10^5^) with a normalized collision energy of 25 eV. In order to avoid repeated sequencing of the same peptides, a dynamic exclusion window of 30 sec was used. In addition, only peptide charge states between two to eight were sequenced.

Raw MS data were processed with MaxQuant software (v1.5.3.30) and searched against the mouse proteome database UniProtKB (UP000000589) with 54,416 entries, released in March 2016. Parameters of MaxQuant database searching were a false discovery rate (FDR) of 0.01 for proteins and peptides, a minimum peptide length of seven amino acids, a first search mass tolerance for peptides of 20 ppm and a main search tolerance of 4.5 ppm. A maximum of two missed cleavages was allowed for the tryptic digest. Cysteine carbamidomethylation was set as fixed modification, while N-terminal acetylation and methionine oxidation were set as variable modifications. The label free quantification (LFQ) algorithm, implemented in MaxQuant, was used for data normalization. The MaxQuant processed output files can be found in the Supporting information, Data S2, showing peptide and protein identification, accession numbers, % sequence coverage of the protein, q-values, and LFQ intensities. The mass spectrometry data have been deposited to the ProteomeXchange Consortium (http://proteomecentral.proteomexchange.org) via the PRIDE partner repository [27] with the dataset identifier PXD017958.

Statistical analysis was done by a two-sample t-test with Benjamini–Hochberg (BH, FDR of 0.05) correction for multiple testing. Significantly regulated proteins between Nf1^Myf5^ and controls were indicated in the Supporting information, Data S3, and were submitted to DAVID bioinformatics 6.8 for functional annotation analysis. For comprehensive proteome data analyses, gene set enrichment analysis (GSEA, v2.2.3) [28] was applied in order to see, if *a priori* defined sets of proteins show statistically significant, concordant differences between Nf1^Myf5^ and controls. GSEA standard settings were used, except that the minimum size exclusion was set to 5 and KEGG v5.2 were used as gene set databases. The cutoff for significantly regulated pathways was set to be ≤ 0.05 p-value and ≤ 0.05 FDR.

### High-Resolution Respirometry

EDL and Soleus muscle from 5 weeks control and Nf1^Myf5^ mice were used in this experiment. Briefly, gently dissected muscles were put into ice-cold biopsy preservation medium (In mM; 2.77 CaK2EGTA, 7.23 K2EGTA, 20 immidazole, 20 taurine, 6.56 MgCl_2_, 5.77 ATP, 3.95 phosphocreatine, 0.5 dithiothreitol, 50 K-MES, pH7.1 at °C) followed by permeabilization with saponin (50μg/ ml) in biopsy preservation medium at 4 °C for 30 min. Fibers were washed with mitochondrial respiration medium (MiR05, 110mM sucrose, 60mM K-lactobionate, 0.5mM EGTA, 3mM MgCl_2_, 20mM taurine, 10mM KH_2_PO_4_, 20mM HEPES, pH 7.1 and 0.1 % fatty acid free BSA) at 4 °C for 10 min. All samples were kept on ice before analysis. The basic principle of this method is to analyze the respiratory capacity at 37 °C in a hyper oxygenated environment with multiple substrate uncouplers. The following concentrations were used: 2mM Malate + 5mM Pyruvate (LEAK respiration), 5mM ADP (OXPHOS capacity), 10μM cytochrome c (integrity of outer mt-membrane), 10mM glutamate (OXPHOS capacity), 10mM succinate (OXPHOS capacity), 0.5μM FCCP (ETS capacity, ETSCI&CII), 0.5μM rotenone (ETS capacity, ETSCII), 2.5μM antimycin A (residual oxygen consumption, ROX) were used in with high-resolution Oxygraph-2k (OROBOROS Instruments). Datlab software (OROBOROS INSTRUMENTS) was used for data acquisition and analysis. Oxygen flux levels were normalized to muscle wet weight of dry blotted fiber bundles.

### Whole Mount in situ hybridization (WMISH)

Embryos at the stage E14.5 were dissected and fixed overnight in 4 % PFA/PBS. Embryos were washed twice in PBST for 30 min on ice and dehydrated through 25 %, 50 % and 75 % methanol in PBST (DEPC-H_2_O), twice in 100 % methanol for 10 min, and were stored in methanol at −20°C. Embryos were rehydrated passing through 75 %, 50 % and 25 % methanol in PBST (DEPC-H2O) and washed twice with 1 x PBS (DEPC-H_2_O) on ice. Afterwards embryos were bleached with 6 % hydrogen peroxide in PBST for 1 h on ice. After washing 3 times for 10 min in PBST, embryos were treated with proteinase K (20μg/ml, Roche) at room temperature for 12 min. Embryos were washed with PBST, PBST/glycine and RIPA buffer for 10 min. each, and fixed with 4% PFA / 0.2% glutaraldehyde for 20 min. followed by washing with PBST, PBST/hybridization buffer and hybridization buffer (50% Formamide, 5x SSC pH 4.5, 1% SDS and 0,1% Tween-20, 0,1 mg/ml tRNA and 0,05 mg/ml Heparin). Embryos were prehybridized in hybridization solution at 65 °C for 3 h, followed by incubation in hybridization buffer containing a Digoxygenin-labelled *MyoD* mRNA probe at 65 °C overnight. Embryos were washed with hybridization buffer at 65 °C for 30 min, followed by digestion with RNaseA at 37 °C for 30 min. Embryos were washed with formamide buffer (50% formamide, 2x SSC pH 4.5, 0,1% Tween-20 in DEPC-water) at 65 °C for 30 min. Followed by two washing steps with MABT for 15 min. each. Embryos were incubated in blocking solution (2% fetal calf serum, 2 mg/ml bovine serum albumin dissolved in TBST) for 1 h and subsequently incubated with anti-DIG-Fab antibody coupled to alkaline phosphatase (Roche: 11093274910) in blocking solution (1 in 5000) at 4°C overnight. Embryos were washed with PBST/tetramisol overnight. Embryos were equilibrated to ALP buffer for three times, 20 min. each, and then transferred to BM purple AP substrate (Roche, 11442074001). For signal conservation the embryos were washed three times with ALP buffer and then fixed in 4 % PFA/PBS. Imaging was performed using a Zeiss Stereo Discovery V12 and the AxioVision 4.6 software (Zeiss). Measurement and quantification of MyoD^+^ stained area was performed with the Autmess function of AxioVision 4.6.

## Results

### Conditional inactivation of Nf1 in myogenic progenitors causes reduction of postnatal muscle growth

To specifically inactivate *Nf1* in the myogenic lineage, we used the Lbx1^Cre^ [22] and Myf5^Cre^ [21] lines, targeting migrating and early committed myogenic progenitors, respectively. Both lines were reanalyzed for their specific activity in the myogenic lineage (Supporting information, *Figure* S1A, B). In the limbs, Lbx1^Cre^ and Myf5^Cre^ were highly specific to the myogenic lineage and did not cause reporter recombination in e.g. neural cells, connective tissue or cartilage. The Myf5^Cre^ line showed minimal off-target activity in dorsal root ganglia, as described previously [21], but no recombination was detected in limb neurons (Supporting information, *Figure* S1A, B). Unlike inactivation of *Nf1* in limb mesenchyme using Prx1^Cre^ [17], inactivation in the myogenic lineage did not cause a marked defect in early myogenesis at embryonal day (E) 14.5 (Supporting information, *Figure* S2A). In late fetal development (E18.5) in both lines, myofiber cross sectional areas were unaltered, while myofiber diameters tended to be decreased, and numbers of central myonuclei were increased (*Figures* 1A – D), indicating a slight developmental delay. Newborn Lbx1^Cre^;Nf1^flox/flox^ mice were smaller, exhibited cyanosis and lack of feeding, and died perinatally. Myf5^Cre^;Nf1^flox/flox^ mice were born at normal mendelian ratios at normal size and were used for further analysis. Myf5^Cre^;Nf1^flox/flox^ mice are further on termed “Nf1^Myf5^”. Myf5^Cre^;Nf1^flox/+^ mice were used as controls unless otherwise noted. Postnatally, Nf1^Myf5^ mice showed a marked growth retardation and weight reduction (*Figures* 1E, F). Dissection of individual muscles (Gastrocnemius, Gas; and Tibialis anterior, TA) showed significantly decreased muscle size (*Figure* 1G) compared to control littermates at seven weeks of age. Efficacy of Nf1 deletion was controlled in E18 and p21 muscle tissue (*Figures* 1H, I). Note that, although the Myf5^Cre^ allele is a loss-of-function, Myf5 mRNA expression in whole muscle tissue both during development (E18.5) and at p21 was indistinguishable between Myf5^+/+^ and Myf5^Cre/+^ mice (Supporting information, *Figure* S2B), indicating compensation by the intact allele. The reduction in muscle size was observed throughout lifespan (*Figure* 1J), which for Nf1^Myf5^ mice consistently was approx. 20 – 25 weeks. Together this suggests a defect in postnatal muscle growth.

**Fig. 1.**
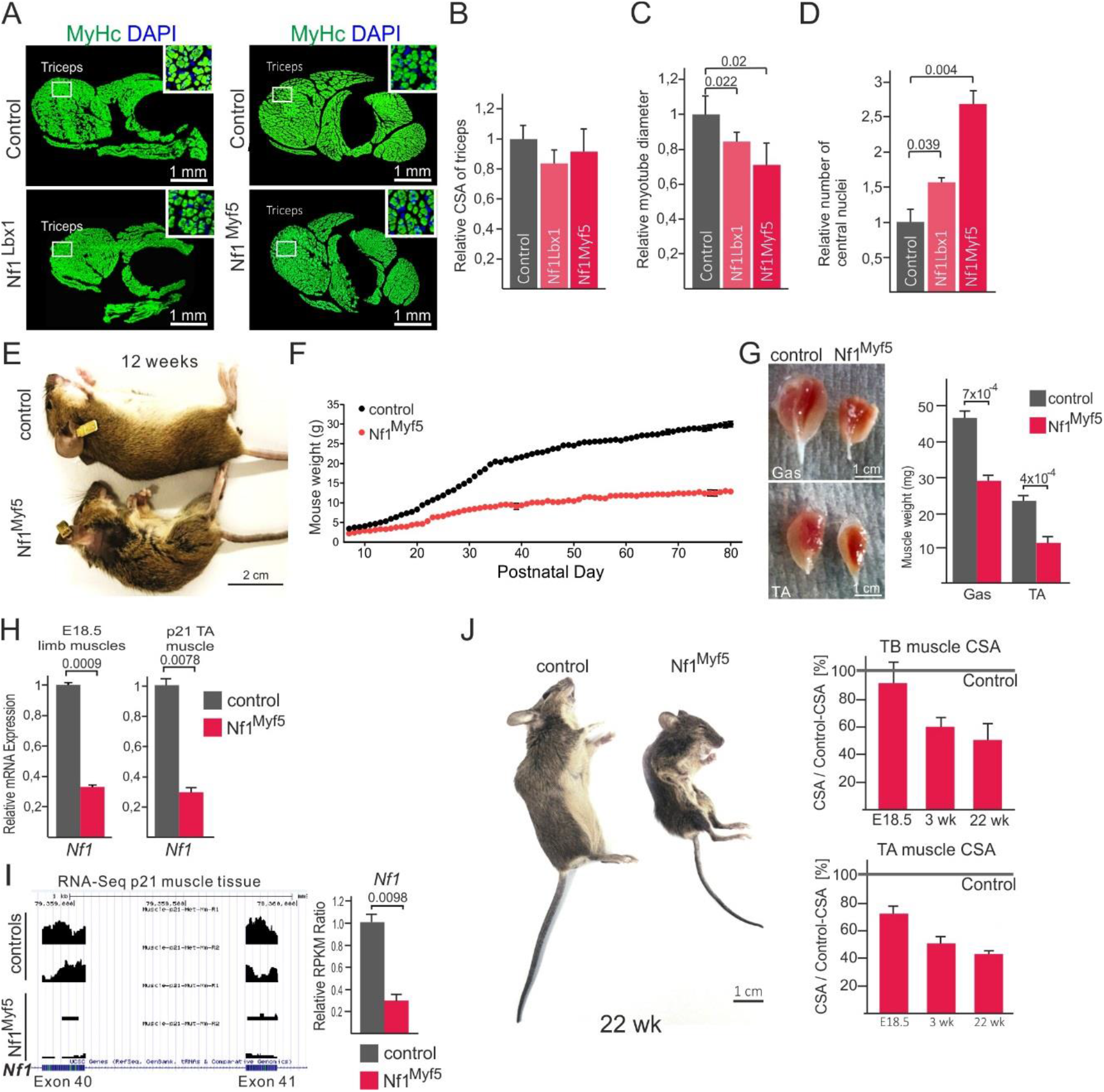
Muscle-specific inactivation of Nf1 causes reduced postnatal muscle growth (A) Forelimb cross sections of control, Nf1^Lbx1^ and Nf1^Myf5^ animals stained for pan-MyHC. Triceps brachii muscle is indicated, squared region is shown as magnification in inserts. (B) Quantification of Triceps brachii cross sectional area (CSA) in TB muscles of controls, Nf1^Lbx1^ and Nf1^Myf5^ animals. (C) Quantification of myotube diameter in TB muscles of controls, Nf1^Lbx1^ and Nf1^Myf5^ animals. (D) Quantification of number of centrally located nuclei in TB muscles of controls, Nf1^Lbx1^ and Nf1^Myf5^ animals. (B – D: n=3 animals per genotype). (E) Control and Nf1^Myf5^ mice at 12 weeks of age. (F) Weight curve of control (black) and Nf1^Myf5^ (red) animals during postnatal life starting at postnatal day 7. (G) Dissected Gastrocnemius (Gas) and Tibialis anterior (TA) muscles of control and Nf1^Myf5^ animals at 7 weeks of age. Quantification is shown right (n=3 animals per genotype). (H) RT-qPCR analysis of *Nf1* deletion efficacy in E18.5 or p21 muscle tissue from control vs. Nf1^Myf5^ mice (n=3 animals per genotype). (I) RNA-Seq tracks showing coverage of *Nf1* exons 40 and 41 in control and Nf1^Myf5^ mice. Quantification is shown right (RPKM: reads per kilobase million). (J) Appearance of 22 week old control and Nf1^Myf5^ animals. Right: Triceps brachii (TB) and Tibialis anterior (TA) cross-sectional area (CSA) of E18.5, p21 and 22 week old Nf1^Myf5^ animals are depicted relative to controls, which was set as 100% (horizontal line) (n=3 animals per genotype).

### Fast fiber atrophy of Nf1^Myf5^ muscles

We next analyzed muscles of young mice amidst the most intense phase of muscle growth, at three weeks of age (p21) by histology. Muscles in fore- and hind limbs (Triceps barchii, TB; Tibialis anterior (TA; Extensor digitorum longus, EDL) showed reduced cross sectional area (*Figures* 2A, B). Centrally located myonuclei were not observed in muscles of p21 or 12 week old Nf1^Myf5^ mice (not shown).

Myofiber numbers were not significantly altered in all three muscles (Supporting information, *Figure* S2C). Conversely, myofiber diameter was reduced in TB, TA and EDL of Nf1^Myf5^ mice (*Figure* 2C). This was confirmed by binning fiber diameters of TB, TA and EDL into discrete size windows, demonstrating global shift towards smaller fibers in Nf1^Myf5^ mice (*Figure* 2D). Staining for type 1 fibers (MyHC-1+) revealed a predominant size reduction of type 2 muscle fibers (*Figure* 2E). We conclude that Nf1^Myf5^ mice show impaired postnatal myofiber growth and persistent fast myofiber atrophy.

**Fig. 2.**
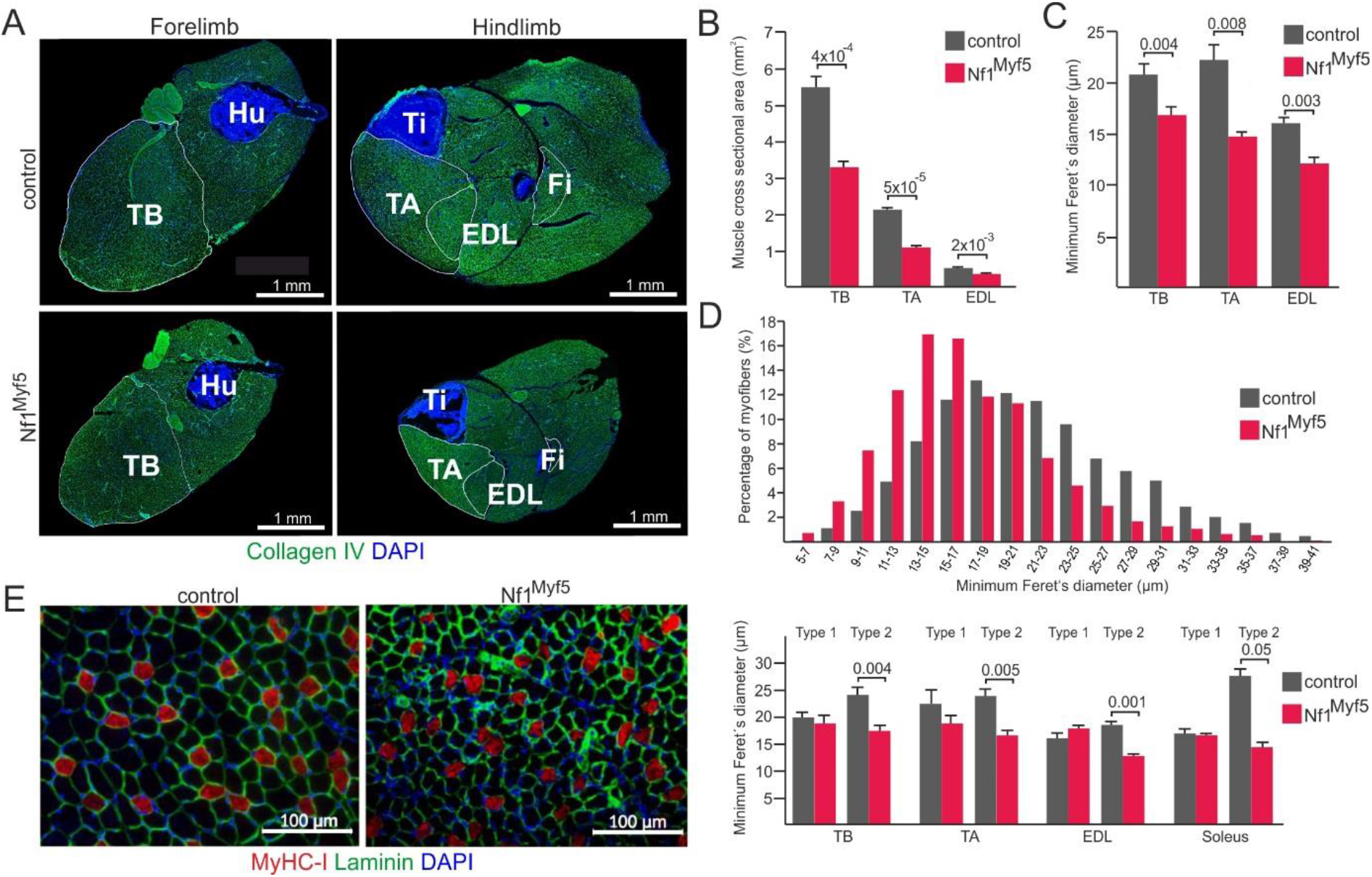
Muscle-specific inactivation of Nf1 causes fast fiber atrophy (A) Cross sections of fore-and hindlimbs of control and Nf1^Myf5^ animals. Staining: Collagen IV (green), DAPI (blue). EDL: Extensor digitorum longus, Fi: fibula, Hu: humerus, TA: Tibialis anterior, TB: Triceps brachii, Ti: tibia. (B) Quantification of the cross sectional area of muscles indicated (n=4 (controls), n=5 (Nf1^Myf5^)). (C) Quantification of minimum Feret’s diameter of individual muscle fibers on tissue cross sections of muscles indicated (n=4 (controls), n=5 (Nf1^Myf5^)). (D) Distribution of myofiber diameter in control and Nf1^Myf5^ animals (n=4 (controls), n=5 (Nf1^Myf5^)). (E) Cross section of TA muscles of control and Nf1^Myf5^ animals. Staining: Myosin heavy chain type 1 (red), Laminin (green) DAPI (blue). Quantification of diameters of type 1 vs type 2 myofibers is shown right (n=3 (controls), n=4 (Nf1^Myf5^)).

### Proteome and transcriptome analysis of Nf1^Myf5^ muscles

To asses global changes in Nf1 deficient muscle tissue we performed proteome as well as transcriptome analysis of whole TA muscles at p21. Note that for the proteome analysis we used both Myf5^Wt^;Nf1^flox/+^ and Myf5^Cre^;Nf1^flox/+^ muscle tissues as controls. Principal component analysis showed that both control genotypes grouped together, while Nf1^Myf5^ samples were grouped separately (Supporting information, *Figure* S3A). This demonstrates that, at least on the proteome level, influence of the Myf5^Cre^ allele (i.e. haploinsufficiency for Myf5), as well as Nf1 haploinsufficiency can be neglected. Proteome analysis of p21 TA muscle lysates yielded 274 significantly deregulated proteins between controls and Nf1^Myf5^ (*Figure* 3A). KEGG pathway analysis showed enrichment for the categories “translation”, “carbon metabolism” and “ribosome” amongst proteins downregulated in Nf1^Myf5^ mutants compared to controls (*Figure* 3B). Amongst proteins upregulated in Nf1^Myf5^ muscle, KEGG terms “oxidative phosphorylation”, “protein digestion”, “proteasome”, “PPAR signaling pathway” and “fatty acid degradation” were overrepresented (*Figure* 3B).

**Fig. 3.**
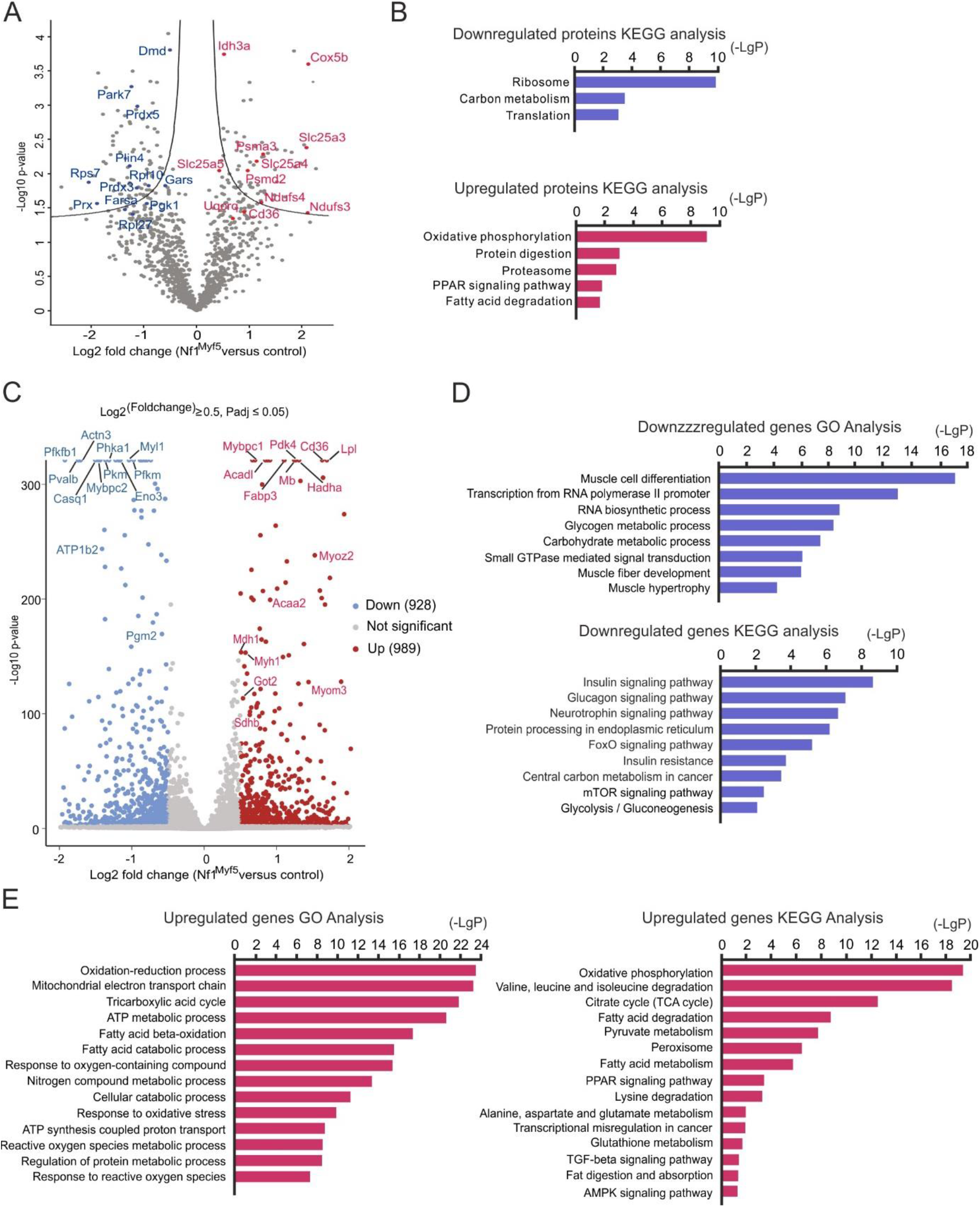
Proteome and transcriptome analysis of postnatal day 21 muscles (A) Volcano plot of control vs. Nf1^Myf5^ muscle proteome data. Individual deregulated proteins are indicated (blue: downregulation, red: upregulation). (B) KEGG pathway analysis of proteins downregulated (top) and proteins upregulated in Nf1^Myf5^ muscle vs. control. (C) Volcano plot of control vs. Nf1^Myf5^ muscle transcriptome data. Individual deregulated transcripts are indicated (blue: downregulation, red: upregulation). (D) Gene ontology (GO) and KEGG analysis of transcripts downregulated in Nf1^Myf5^ muscle vs. control. (E) GO and KEGG analysis of transcripts upregulated in Nf1^Myf5^ muscle vs. control.

This was confirmed by RNA-Seq analysis of p21 TA muscle tissue. Principal component analysis confirmed separate clustering of control and Nf1^Myf5^ samples (Supporting information, *Figure* S3B). Transcriptome analysis yielded 1917 differentially expressed genes; 928 genes were down-, 989 genes were upregulated in Nf1^Myf5^ muscle relative to controls (*Figure* 3C). GO as well as KEGG analysis of transcriptome data highlighted terms as “muscle cell differentiation”, “muscle hypertrophy”, “carbohydrate metabolism” or “glycolysis/gluconeogenesis” as overrepresented in genes downregulated in Nf1^Myf5^ muscle (*Figure* 3D). Conversely, terms as “oxidation-reduction process”, “mitochondrial electron transport chain”, “TCA cycle” or “oxidative phosphorylation”, as well as several terms related to amino acid catabolism were enriched in genes upregulated in Nf1^Myf5^ muscle (*Figure* 3E). We also noted “insulin signaling pathway” enriched in downregulated genes; we therefore first analyzed Nf1^Myf5^ mice for blood glucose levels, which were unchanged (Supporting information, *Figure* S4A). Also, phosphorylation of Akt as well as IRS1 in random fed animals were not significantly changed (Supporting information, *Figure* S4B). In summary, proteome and transcriptome analysis suggest a) metabolic shift and b) deranged protein homeostasis in Nf1^Myf5^ mice.

### Increased oxidative metabolism in Nf1^Myf5^ muscle

Skeletal muscle primarily gains energy by breakdown of carbohydrates and fatty acids, both being able to feed into mitochondrial respiration as the most efficient way of ATP production. In Nf1^Myf5^ muscle, GO as well as KEGG categories associated with the TCA cycle or the mitochondrial electron transport chain were overrepresented in genes and proteins upregulated in Nf1^Myf5^ muscle (*Figure* 3B, E). Unbiased gene set enrichment analysis (GSEA) of the RNA-Seq data highlighted enrichment of genes associated with “glucose catabolic process” in control, and of genes associated with “oxidative phosphorylation” in Nf1^Myf5^ muscle (*Figure* 4A). We therefore mined the differentially expressed gene set for genes encoding components of glycolysis or TCA cycle / oxidative phosphorylation, respectively. This showed that glycolysis appeared globally downregulated, with exception of *Pdk2, Pdk4* and *Ldhb,* while TCA cycle and OXPHOS genes were globally upregulated (*Figure* 4B, C). In line, detailed analysis of proteome data showed several proteins of the TCA cycle as well as the electron transport chain upregulated in Nf1^Myf5^ muscle (*Figure* 4D).

**Fig. 4.**
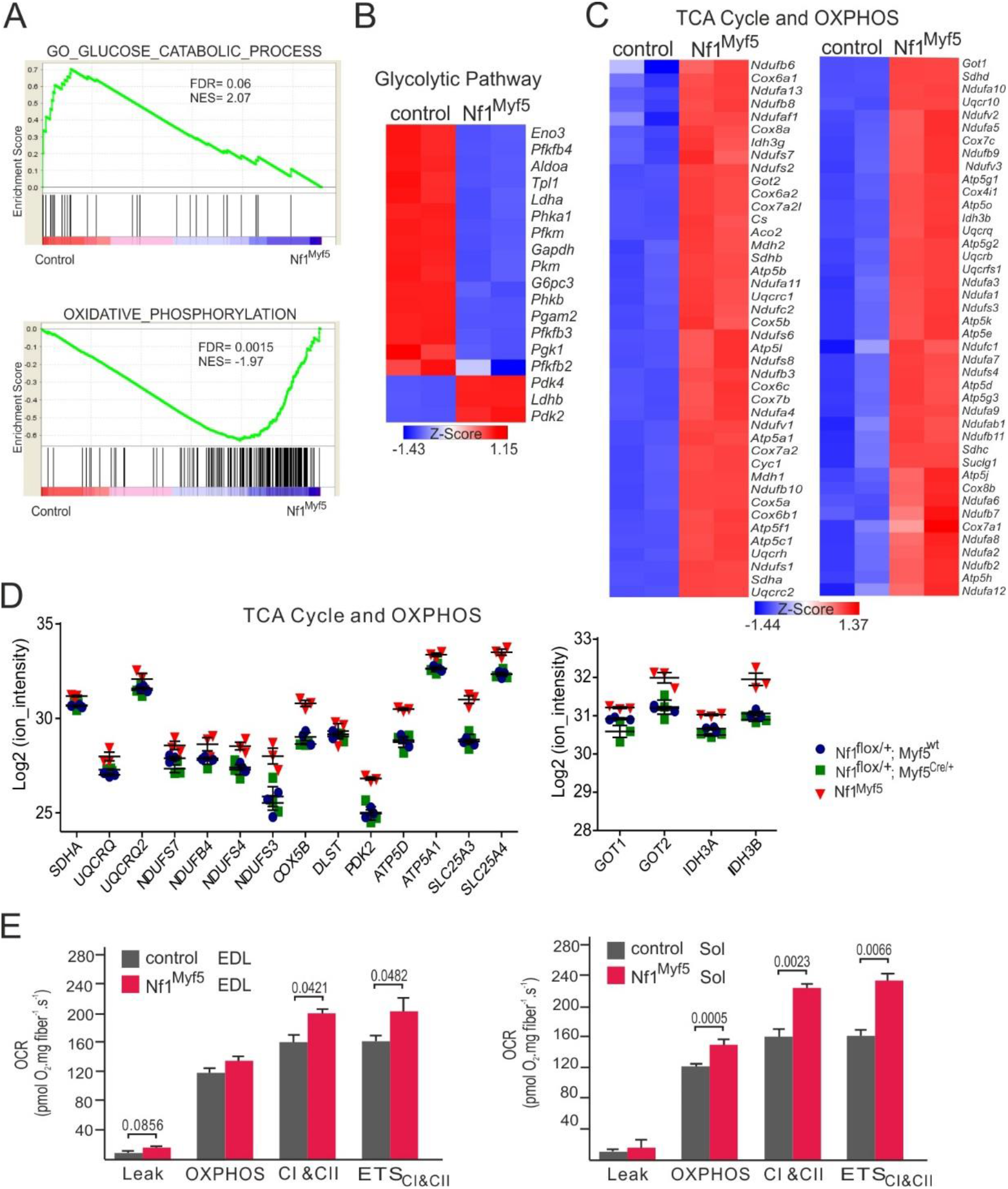
Disrupted metabolic homeostasis in Nf1^Myf5^ muscle (A) Gene set enrichment analysis (GSEA) of raw control vs. Nf1^Myf5^ muscle transcriptome data for the terms “glucose catabolic process” and oxidative phosphorylation”. (B) Heatmap of differentially expressed glycolytic genes (control vs. Nf1^Myf5^ muscle). (C) Heatmap of differentially expressed genes (control vs. Nf1^Myf5^ muscle) of tricarboxic acid (TCA) cycle and oxidative phosphorylation machinery (OXPHOS) components. (D) Depiction of significantly deregulated proteins related to TCA cycle and OXPHOS. Proteome analysis of p21 muscle, genotypes are indicated. (E) Real-time respirometry measurement of control vs. Nf1^Myf5^ EDL (Extensor digitorum longus) and Sol (Soleus) muscles (n=7 animals for both genotypes).

To directly measure mitochondrial oxidative metabolism, we employed high-resolution respirometry on freshly isolated muscle fibers from 5 weeks old control and Nf1^Myf5^ mice. We analyzed the Extensor digitorum longus (EDL) muscle as a prototypical prevalently fast glycolytic muscle, and the Soleus muscle as a prototypical oxidative muscle for comparison. This clearly demonstrated increased mitochondrial respiratory capacity in muscles of Nf1^Myf5^ animals (*Figure* 4E).

### Glycolytic-to-oxidative fiber type shift in Nf1^Myf5^ muscle

Most muscles of the limbs consist of predominantly fast-twitching type 2 fibers, which are larger in diameter than slow-twitching type 1 fibers and produce main mechanical force. Type 2 fibers can be subdivided into Type 2B fibers that show anaerobic glycolytic metabolism, and type 2A fibers that use aerobic metabolism. Type 2B fibers express Myosin heavy chain type 2B (MyHC-2B) or a combination of MyHC-2B and MyHC-2X (encoded by the *Myh4* and *Myh1* genes, respectively). Type 2A fibers express MyHC-2A encoded by *Myh2,* and type 1 fibers express MyHC-1 encoded by *Myh7.* Mining the p21 transcriptome data for markers of fast-vs. slow fibers showed reduction of fast markers and increased expression of slow markers in Nf1^Myf5^ muscles (*Figure* 5A). Quantitative real-time PCR (RT-qPCR) confirmed upregulation of *Myh2* and *Myh7,* and downregulation of *Myh4* mRNA in Nf1^Myf5^ muscle (*Figure* 5B). In accordance, protein levels of MyHC-1 were increased, and levels of MyHC-2B were decreased in Nf1^Myf5^ TA muscles (*Figure* 5C). This indicated distorted fast / slow fiber composition in Nf1^Myf5^ muscles.

**Fig. 5.**
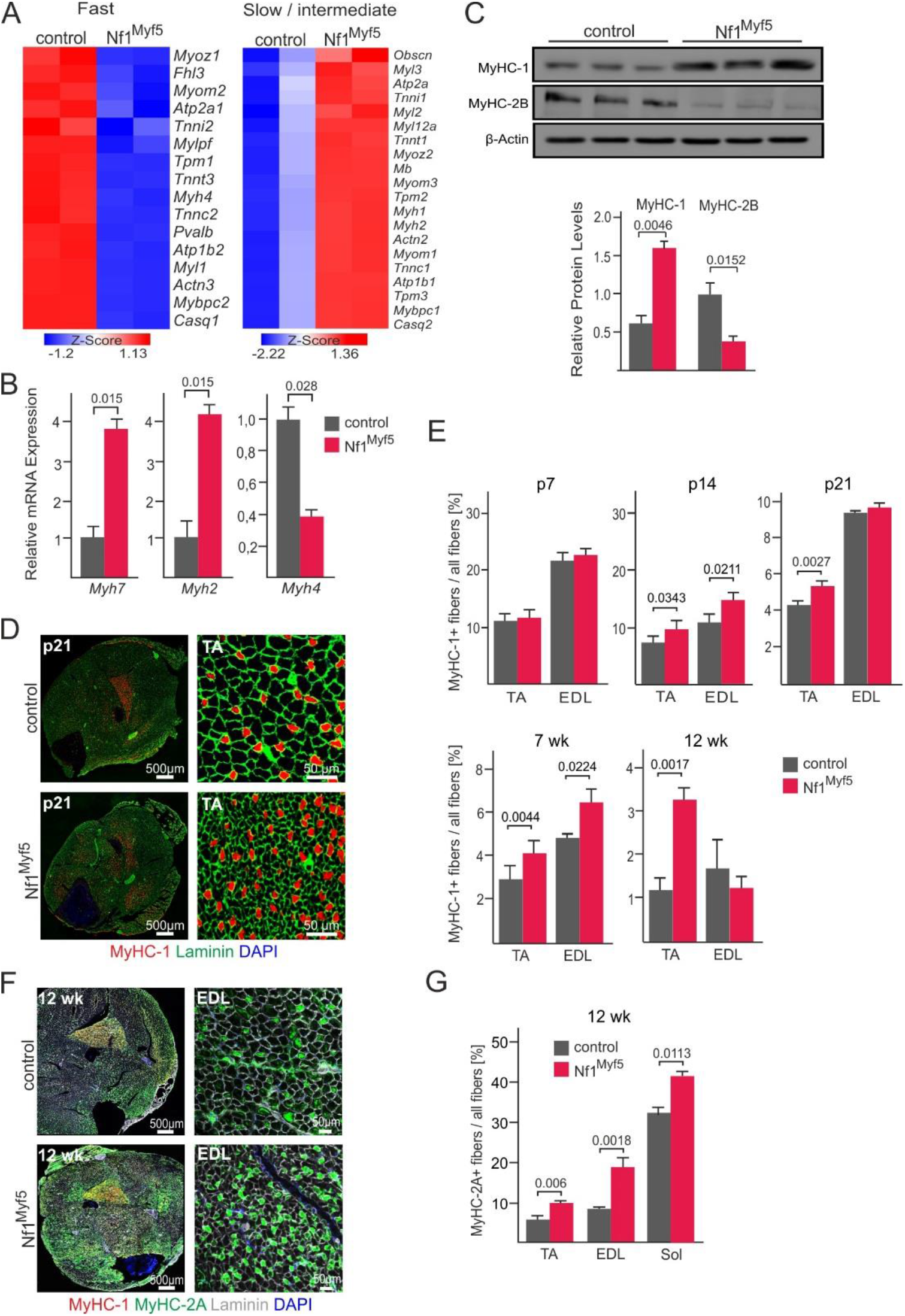
Fiber type shift in Nf1^Myf5^ muscle (A) Heatmaps showing differentially expressed genes specific for fast or slow / intermediate fibers, respectively between control vs. Nf1^Myf5^ muscle. (B) RT-qPCR analysis of Myosin heavy chain genes *Myh7, Myh2* and *Myh4* on control vs. Nf1^Myf5^ muscle (n=3 animals per genotype). (C) Western Blot analysis of myosin heavy chain type 1 (MyHC-1) and type 2B (MyHC-2B) on control vs. Nf1^Myf5^ muscle. Quantification below (n=3 animals per genotype). (D) Immunolabeling for MyHC-1 on cross sections of p21 control vs. Nf1^Myf5^ lower legs. TA muscles are shown as magnifications right. Sections are stained for Laminin and DAPI. (E) Quantification of Type 1 fibers / all fibers for indicated postnatal time points on TA and EDL muscles (n=3 animals per genotype). (F) Immunolabeling for fast oxidative fiber type myosin (MyHC-2A) and MyHC-1 on cross sections of control vs. Nf1^Myf5^ lower legs. Sections are stained for Laminin and DAPI. EDL muscles are shown as magnifications right. (G) Quantification of percentage of MyHC-2A fibers / all fibers (n=3 animals per genotype).

The expression of fast-fiber related genes is induced in the first weeks of postnatal life in rodents [13,29]. We therefore analyzed the number of type 1 fibers in TA and EDL muscles (which are predominantly composed of fast fibers) over time (*Figure* 5D, E). At postnatal day 7 (p7), no differences were observed. But from postnatal day 14 (p14) on a persistent slight increase in type 1 fibers in the TA muscle, and a transient increase in the EDL muscle in Nf1^Myf5^ mice during postnatal development were seen (*Figures* 5D, E). We then analyzed the proportion of type 2A and type 2B fibers in adult mice, revealing a shift of type 2B/X to type 2A fibers, i.e. towards the fast oxidative phenotype, in Nf1^Myf5^ mice (*Figures* 5F, G). We conclude that Nf1^Myf5^ muscles show a metabolic shift towards oxidative metabolism with partial conversion of type 2B/X to type 2A or type 1 fibers.

### Defective protein homeostasis in Nf1-deficient muscles

Proteome analysis showed a significant downregulation of several components of the translational machinery, especially ribosomal proteins as well as tRNA synthases (*Figures* 3A, 6A). Conversely, several proteasomal subunits were upregulated (*Figures* 3A, 6B). This may indicate decreased protein synthesis and increased breakdown suggesting a mechanism paralleling myofiber atrophy. In line, RT-qPCR analysis demonstrated upregulation of common atrophy-related transcripts [30–32] in Nf1^Myf5^ muscle (*Figure* 6C). These genes comprise *Ctsl* (Cathepsin L) and *Psma1* (Proteasomal subunit alpha 1), or the classical atrogenes Fbxo32 (Atrogin-1 / MAFbx) and Trim63 (MuRF1) [33]. Myofiber atrophy typically is caused by a dysbalance between protein synthesis and breakdown. Protein synthesis in skeletal muscle is mainly regulated via mTORC1 [11,12]. Interestingly, the KEGG term “mTOR signaling pathway” was enriched in genes downregulated in Nf1^Myf5^ muscle (*Figure* 3D). To analyze mTORC1 activity we assessed phosphorylation of mTOR as well as the mTORC1 downstream target S6 ribosomal protein. p21 Nf1^Myf5^ muscle tissue showed hallmarks of decreased mTORC1 signaling, with decreased phosphorylation of mTOR Ser-2448, and decreased S6 Ser-235/236 phosphorylation (*Figure* 6D). Altogether this suggests disturbed protein homeostasis in Nf1^Myf5^ muscle with decreased mTORC1-driven protein synthesis and increased protein breakdown. Such alterations may be the result of a muscular energy deficit [11]. In line, Nf1^Myf5^ muscle showed increased levels of phosphorylated AMP-dependent kinase (AMPK), the most prominent intracellular energy sensor (*Figure* 6E). In sum, Nf1^Myf5^ animals show typical signs of muscle atrophy with decreased anabolic signaling, in combination with an unmet energy demand.

**Fig. 6.**
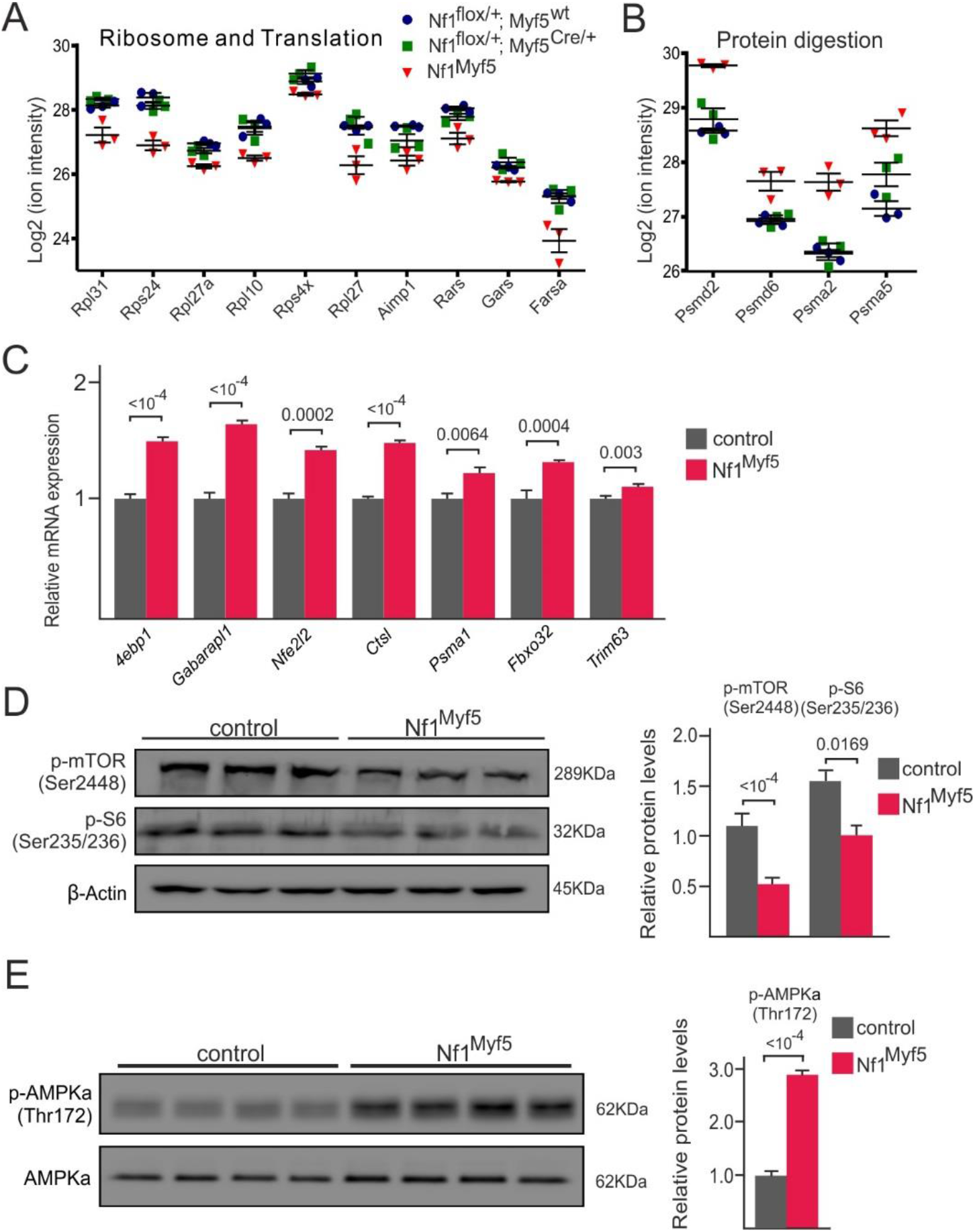
Disturbed protein homeostasis and energy deficit in Nf1^Myf5^ muscle (A) Depiction of significantly deregulated components of the ribosome and proteins related to translation. Proteome analysis of p21 muscle, genotypes are indicated. (B) Depiction of significantly deregulated proteins related to protein digestion. Proteome analysis of p21 muscle, genotypes are indicated. (C) RT-qPCR analysis of p21 control and Nf1^Myf5^ muscle for transcripts related to muscle atrophy (n=3 animals per genotype). (D) Western blot analysis for phosphorylated mTOR (Ser2448) and phosphorylated S6 ribosomal protein (Ser235/236): Quantification shown right (n=3 animals per genotype). (E) Western blot analysis for phosphorylated AMPK (Thr172). Quantification shown right (n=4 animals per genotype).

### Increased fatty acid catabolism in muscle and paucity of white adipose tissue in Nf1^Myf5^ mice

Increased oxidative metabolism, but concomitant downregulation of glycolytic genes indicated that Nf1^Myf5^ muscles may predominantly use fatty acids to feed the TCA cycle. Analysis of proteome as well as transcriptome data highlighted terms associated with fatty acid catabolism, as “fatty acid degradation” (proteome) or “fatty acid beta-oxidation” (transcriptome) were significantly enriched amongst proteins / mRNAs upregulated in Nf1^Myf5^ muscle (*Figure* 3B, E). Accordingly, several proteins associated with fat digestion and absorption were upregulated in Nf1^Myf5^ muscle (*Figure* 7A). In line with GO and KEGG analysis of differentially expressed genes, unbiased GSEA of transcriptome data showed genes belonging to the clusters “fatty acid beta-oxidation” and “fatty acid catabolic process” strongly enriched in Nf1^Myf5^ muscle (*Figure* 7B). Indeed, all genes belonging to the GO term “fatty acid metabolism” were upregulated in Nf1^Myf5^ muscle (*Figure* 7C), including e.g. the gene encoding the main fatty acid transporter CD36, or several Acyl-CoA-dehydrogenases, which not only play a role in β-oxidation but also breakdown of branched amino acids. Upregulation of several genes of this panel in Nf1^Myf5^ muscle (*Lpl* encoding part of the enzyme complex responsible for lipid uptake, Fatty acid binding proteins *Fabp3* and *Fabp4,* Carnithine palmitoyltransferases *Cpt1b* and *Cpt2,* and *Acad1* encoding Medium chain acyl-CoA dehydrogenase) were confirmed by RT-qPCR (*Figure* 7D). In sum, proteome as well as transcriptome data indicate increased fatty acid breakdown in Nf1^Myf5^ muscle to fuel the TCA cycle and OXPHOS. In line with this, and contrasting previous reports [20], we did not find indication of ectopic lipid storage in muscle fibers in our muscle-specific Nf1 mutants at 1 or 12 weeks of age (*Figure* 7E; see supporting information, *Figure* S5A for overview images).

**Fig. 7.**
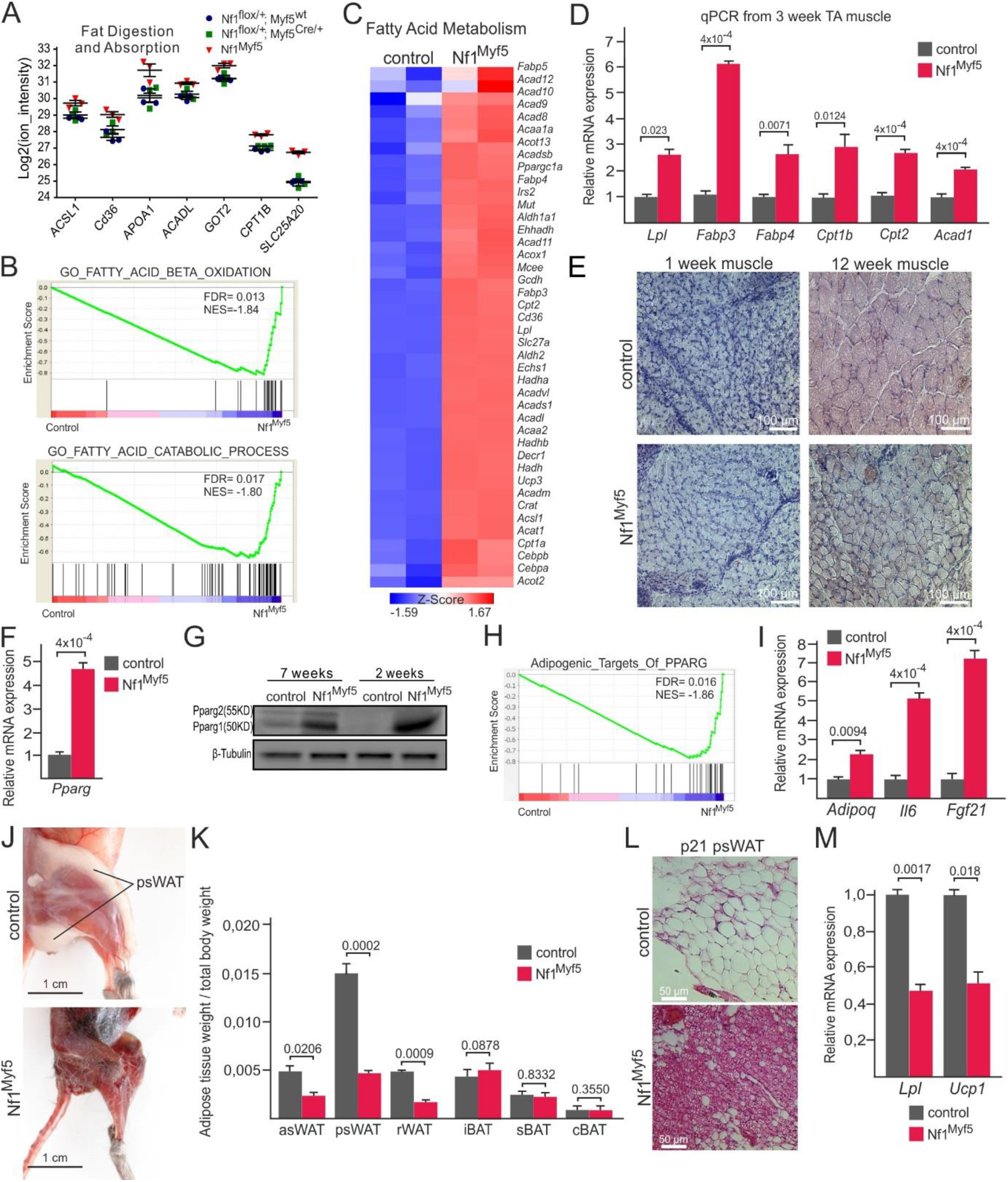
Increased fatty acid oxidation in Nf1^Myf5^ muscle (A) Depiction of significantly deregulated proteins related to fat digestion and absorption. Proteome analysis of p21 muscle, genotypes are indicated. (B) Gene set enrichment analysis (GSEA) of raw control vs. Nf1^Myf5^ muscle transcriptome data for the terms “fatty acid beta oxidation” and fatty acid catabolic process”. (C) Heatmap of differentially expressed genes related to fatty acid metabolism (control vs. Nf1^Myf5^ muscle). (D) RT-qPCR analysis in control vs. Nf1^Myf5^ muscle of selected genes related to fatty acid uptake, transport and catabolism (n=3 animals per genotype). (E) Oil red O lipid staining on control vs. Nf1^Myf5^ muscle (1 week, 12 weeks). (F) RT-qPCR analysis of *Pparg* in control vs. Nf1^Myf5^ muscle (n=3 animals per genotype. (G) Western blot analysis for PPARγ on control vs. Nf1^Myf5^ muscle. (H) GSEA of control vs. Nf1^Myf5^ muscle transcriptome data for the term “adipogenic targets of PPARG”. (I) RT-qPCR analysis for *Adipoq, Il6* and *Fgf21* on control vs. Nf1^Myf5^ muscle (n=3 animals per genotype). (J) Nf1^Myf5^ (p21) mice show reduced posterior subcutaneous white adipose tissue (WAT). Note dark red appearance of muscles in Nf1+ mutant. (K) Weight of adipose tissue depots of control vs. Nf1^Myf5^ mice at 12 weeks of age. Adipose depot weight was normalized to body weight. AsWAT: anterior subcutaneous white adipose tissue; psWAT: posterior subcutaneous white adipose tissue; rWAT: retroperitoneal white adipose tissue; iBAT: interscapular brown adipose tissue; sBAT: subscapular brown adipose tissue; cBAT: cervical brown adipose tissue (n=4 animals per genotype). (L) Histological analysis (hematoxylin and eosin) of p21 psWAT of control vs. Nf1^Myf5^ mice. (M) RT-qPCR analysis of p21 psWAT of control vs. Nf1^Myf5^ mice for *Lpl,* and *Ucp1* (n=3 animals per genotype).

PPARγ has been involved in promoting intramuscular lipolysis [34,35]. In muscle tissue of Nf1^Myf5^ mice we found overexpression of *Pparg* mRNA (*Figure* 7F); specific upregulation of PPARγ isoform 1 was confirmed on the protein level in 7 week and also already in two week old animals (*Figure* 7G). In line, GSEA showed enrichment for “Adipogenic targets of PPARG” enriched in Nf1 mutants (*Figure* 7H). Similarly, “PPAR signaling pathway” was enriched in transcripts and proteins upregulated in Nf1 mutants (*Figures* 3B, E). PPARγ was shown to induce the myokine Adiponectin (Adipoq) [36]. Nf1 deficient muscle showed increased expression of *Adipoq* as well as mRNAs of the myokine *Il6* and the catabolic cytokine *Fgf21* (*Figure* 7I).

Finally, increased fatty acid consumption in Nf1^Myf5^ muscles as indicated by proteome and transcriptome analysis suggests increased lipid mobilization from adipose tissue. Indeed, p21 Nf1 mutants showed a drastic reduction of their white adipose tissue (WAT), affecting both subcutaneous as well as visceral WAT (*Figures* 7J, K). This was specific to WAT, brown adipose tissue (BAT) depots were unaffected (*Figure* 7K). In histology, WAT appeared denser, deprived of lipid content (*Figure* 7L). RT-qPCR analysis showed decreased expression of *Lpl* in WAT of Nf1^Myf5^ mice (*Figure* 7M) suggesting decreased lipid uptake. *Ucp1* was not induced, but rather downregulated in Nf1^Myf5^ WAT (*Figure* 7M), suggesting no “browning” of WAT. Expression of *Pparg* was unaltered in Nf1^Myf5^ psWAT, as were *mtCo1* and *Cs* (encoding Mitochondrial cytochrome c oxidase and Citrate synthase, respectively) as representatives of electron transport chain and TCA cycle (Supporting information, *Figure* S5B). BAT originates in large part from a dermomyotomal Myf5+ progenitor population [37], thus Nf1 inactivation in BAT may influence the phenotype. Intriguingly, RT-qPCR analysis of BAT showed some overlap in gene deregulation with Nf1-deficient muscle, as *mtCo1* was upregulated, however *Cs*, *Pparg,* and also *Ucp1* were unaltered (Supporting information, *Figure* S5C) suggesting no major influence on BAT. This altogether suggests a general catabolic state of Nf1^Myf5^ animals forced by unmet energetic need of muscle tissue leading to a cachectic state with increased lipid mobilization.

## Discussion

We show here that Nf1 inactivation specifically in the myogenic lineage leads to postnatal myopathy mirroring a key feature seen in Nf1 patients. Thus, the Nf1^Myf5^ line enables analysis of Nf1 function in postnatal muscles, and in comparison to models as the Nf1^Prx1^ mouse is suitable to distinguish muscle-specific from non-myogenic functions of Nf1. Nf1^Myf5^ mice are viable, while Nf1^Lbx1^ (this study) and Nf1^Myod^ mice [20] were postnatally lethal. This difference may be caused by different efficacy of the Cre driver lines used. Lbx1^Cre^ efficiently targets early migrating myogenic progenitors, including those of the diaphragm and tongue [22] and expression of *Lbx1* precedes the expression of *Myf5* [38]. While MyoD is expressed in all myogenic cells, not all myoblasts have experienced expression of Myf5, although there is dispute whether Myf5 and Myod cells represent partially separate lineages [39–42]. Thus, very likely not all myogenic cells are targeted in the Nf1^Myf5^ model, possibly alleviating the phenotype to a life-compatible level. Depletion of the Myf5 lineage led to compensatory takeover by Myod-lineage cells [39], and Comai et al. [40] demonstrated a marked compensatory effect of recombination escapers upon Myf5 cell depletion. Importantly, robust knockdown of *Nf1* mRNA in Nf1^Myf5^ mice at p21 argues against a possible lineage takeover of Myf5-negative cells or of recombination escapers, suggesting no significant skewing of the myogenic lineage had occurred.

Nf1^Myf5^ mice did not show increased interstitial fibrotic and adipogenic infiltration of muscle as it was seen after inactivation of Nf1 in limb mesenchyme via Prx1^Cre^ [17]. Muscle interstitial fibroblasts and adipocytes do not originate from myogenic cells, but from non-myogenic mesenchymal progenitors, the so-called fibro-adipogenic progenitors [43–45]. These cells in turn derive from lateral plate mesoderm-derived limb bud mesenchyme [46]. The same developmental mesenchymal progenitors also form muscle interstitial fibroblasts and adipocytes during development [47]. Prx1^Cre^ targets the entire limb mesenchyme [19] encompassing this mesenchymal progenitor population during development [48]. Thus, comparing Nf1^Prx1^ and Nf1^Myf5^ models, we conclude that fibrofatty infiltration of muscle in the NF1^Prx1^ model does not originate from myogenic cells but from interstitial mesenchymal cells. However, in another model of muscle-specific inactivation of Nf1 (Myod^Cre^) ectopic intramyocellular lipid accumulation was reported [20], which we did not observe in our model. This discrepancy might be caused by the different Cre drivers used; of note, Myod^Cre^ was reported to be active in the developing liver [49] thus an indirect effect may not be excluded.

Nf1^Myf5^ mice show reduced postnatal fiber growth that manifested early in life. This suggests that loss of Nf1 does not induce muscle atrophy at a specific time point, but rather that NF1-associated muscle weakness is a continuous process of decreased hypertrophic growth beginning in childhood. Indeed, muscle weakness and decreased muscle CSA are already seen in pediatric patients [7,8]. In Nf1^Myf5^ mice predominantly fast (type 2) fibers are affected. In general, type 2 fibers have higher potential for hypertrophy than type 1 fibers, which also makes them more vulnerable for atrophy [50]. Nevertheless, different myopathies predominantly affect different fiber types, the underlying cause is mostly unclear [14]. Our results indicate dysbalanced protein homeostasis in Nf1^Myf5^ muscle. TORC1 signaling is a master regulator for protein synthesis [11,12]. Nf1^Myf5^ muscle showed reduced mTORC1 activation that is in line with proteome and transcriptome data, altogether indicating decreased capacity for protein synthesis. On the other hand common atrogenes were upregulated on the transcript level, and proteasomal components were upregulated on transcript and protein level, in addition suggesting increased protein breakdown.

In parallel we observed a fiber type shift from glycolytic type 2 fibers to oxidative type 2 and in part to type 1 fibers. In adult Nf1^Prx1^ mice, no fiber type shift was reported [18], however this had not been quantified. During embryonic and fetal myogenesis, muscle fibers predominantly show oxidative-type gene and protein expression. Fast-type gene and protein expression is first detected during late fetal development, the major slow-to-fast fiber type change occurs during the early postnatal period [13]. The emergence of different fiber types in development is under control of transcription factors such as *Six1, Six4, Sox6, Tbx15, Prdm1* (aka *Blimp1)* and *Nfix* [51–57], however we did not detect a significant deregulation of these factors in our transcriptome data (Supporting information, Data S1). Mature fibers have a degree of plasticity and can shift their metabolism based on e.g. endurance vs. peak force training [13]. Importantly, fiber type shifts are involved in muscular disorders and muscle aging [13,14]. Nf1 is best known as a RAS-GAP that negatively regulates downstream MEK-ERK signaling [1]. Interestingly, ERK1/2 activity itself was involved in regulating muscle fiber types previously [13]. In line it was recently shown that expression of an activated form of MEK1 in muscle fibers can drive a metabolic shift to oxidative fibers [58].

In line with increased oxidative metabolism relying on fatty acid consumption, Nf1^Myf5^ mice showed severe reduction of WAT. Myf5^Cre^ mediated gene inactivation preferentially targets the myogenic lineage, however it also targets brown adipose tissue as well as several (anteriorly located) white adipose depots [59]. Thus, inactivation of *Nf1* in subsets of adipocyte progenitors may have a direct effect on adipogenesis / adipocyte function. Inactivation of the tumor suppressor *Pten,* leading to hyperactivation of the PI3K pathway, in the Myf5 lineage led to increased size of anterior BAT and WAT depots, while posterior WAT depots were virtually absent in mutants [60]. We did not detect differences in BAT size in Nf1^Myf5^ mice. Moreover, *Ucp1* mRNA was not upregulated in BAT, indicating that Nf1-deficient BAT does not show increased energy expenditure. WAT depots were reduced irrespective of anatomical location; asWAT and rWAT that are targeted by Myf5^Cre^, as well as psWAT that is not targeted [59] were strongly affected. This argues against a cell autonomous effect on *Nf1* deletion in adipogenic progenitors / adipocytes, but rather in favor of a muscle-derived effect enforcing lipolysis in WAT. We observed upregulation of mRNAs encoding myokines, i.e. muscle-produced endocrine signaling factors. FGF21 and IL6 were shown to stimulate lipolysis in WAT [61,62]. IL6 also stimulates lipolysis and fatty acid oxidation in muscle in an autocrine fashion, interestingly via AMPK activation [63–65]. Furthermore, *Adipoq,* encoding adiponectin, was induced. Adiponectin is mainly known as an adipokine, however also skeletal muscle produces Adiponectin, and *Adipoq* KO mice show increased intramyocellular lipid content [66]. Adiponectin expression in muscle can be induced by PPARγ [36], which was overexpressed in Nf1^Myf5^ muscle. PPARγ in muscle promotes fatty acid oxidation [34,35]. PPARγ overexpression in muscle protects from obesity-induced diabetes, leads to increased adiponectin expression, induces oxidative metabolism, and muscles have reduced lipid content [36]. Conversely, muscle specific inactivation of PPARγ induces obesity and insulin resistance [67,68]. FGF21 is well described as mitochondrial stress-induced myokine [69–71]. Very recently, FGF21 was described as a novel player in the regulation of muscle mass during fasting-induced muscle atrophy and weakness [72]. Here, we describe a strong induction of muscle *Fgf21* in response to genetic ablation of Nf1, together with a dramatic reduction of WAT depots, affecting both subcutaneous as well as visceral WAT. However, future studies are warranted to clarify the protective or detrimental catabolic action of FGF21 via inter-tissue crosstalk in Nf1^Myf5^ animals.

Contrasting increased oxidative metabolism and fatty acid breakdown, downregulation of all glycolysis components in the transcriptome analysis indicated decreased glycolytic flux in Nf1^Myf5^ muscles. Intriguingly, *Pdk2, Pdk4* and *Ldhb* were upregulated. The B-isoform of LDH is mainly expressed in oxidative muscle, and overexpression of *Ldhb* increases oxidative metabolism [73] in line with the oxidative phenotype we observe. PDK2 and PDK4 negatively regulate Pyruvate dehydrogenase and thus impair funneling Pyruvate from glycolysis towards usage in the TCA cycle, shunting mitochondrial metabolism towards fatty acid usage [13]. This altogether suggests that utilization of glucose as fuel for the TCA cycle is impaired in Nf1^Myf5^ muscle, thus fatty acids as alternative energy source need to be used. Intriguingly, a decreased respiratory quotient indicating lower carbohydrate and increased lipid oxidation was demonstrated in NF1 patients [74]. Thus defective glycolysis may therefore explain why Nf1^Myf5^ muscles shift towards oxidative metabolism and primarily rely on fatty acids, but could also contribute to the apparent energy deficit of Nf1-deficient muscle indicated by AMPK activation. Nf1 is well-known as a tumor suppressor gene repressing RAS signaling. Increased RAS signaling or loss of Nf1 is typically associated with the induction of a Warburg effect, i.e. increased glycolytic flux under aerobic conditions, as well as the induction of pathways supporting cell growth, as mTORC1 [75,76]. Why loss of Nf1 in myogenic cells leads to effectively opposite outcome is unknown, however it indicates differences in either the cellular environment or intracellular signal reception that lead to a different interpretation of Nf1-dependent signals. Decreased energy charge is sensed in cells via AMPK that is a master regulator of cellular energy homeostasis [77]. Activation of AMPK can reduce mTORC1 activity and thus protein synthesis, providing a possible explanation for the reduced hypertrophic growth of Nf1^Myf5^ muscles. In turn, AMPK activation is known to induce mitochondrial biogenesis, glucose uptake and fatty acid consumption [77,78]. This altogether suggests that impaired glycolysis may be the primary defect in Nf1^Myf5^ muscle, leading to an energy deficit, shifting cellular metabolism via AMPK towards OXPHOS.

Mitochondria are described as the predominant source of reactive oxidant species (ROS) in muscle fibers [79]. Notably, oxidative stress in skeletal muscles can promote protein synthesis/degradation imbalance and muscle wasting [80]. Increased ROS levels in cachectic muscle could be attributed to the reduction in the activity of endogenous antioxidant enzymes [81]. In drosophila, loss of Nf1 caused mitochondrial oxidative stress concomitant to decreased ATP production [82]. GO and KEGG analysis of transcriptome data showed enrichment of the terms “Response to oxygen-containing compound, “Response to oxidative stress”, “Reactive oxygen species metabolic process”, “Response to reactive oxygen species (GO), and “Glutathione metabolism” (KEGG) in genes upregulated in Nf1^Myf5^ muscle (Fig. 3E). In addition transcriptome analysis showed increased expression of NADPH oxidase genes *Cybb (Nox2)* and *Nox4* (Supporting information, Data S1), which are an additional source of ROS [83]. This indicates that increased mitochondrial respiration in Nf1^Myf5^ muscle may lead to oxidative stress, hampering mitochondrial energy production. This raises the question whether a concomitant increased mitochondrial ROS production may even aggravate the situation in Nf1^Myf5^ mice, potentially causing a vicious cycle.

In summary, Nf1^Myf5^ represents a viable mouse model for muscle-specific inactivation of Neurofibromin that recapitulates the pediatric myopathy seen in NF1 patients. Nf1 deficiency disrupts metabolic homeostasis in muscle leading to muscular energy deficit and a catabolic state causing reduced postnatal muscle hypertrophy.

## Author Contributions

X.W., and S.S. developed the study concept and design. X.W. performed the majority of experiments and data collection. J.G., M.O., K.W., A.C.P., S.B., B.T., D.M., and S.K., performed additional experiments or provided resources. X.W., M.O., D.M., A.K., and S. S. interpreted data. X. W., and S.S. prepared the figures. M.O., A.K., S.K., and S. S. wrote the manuscript. All authors declare that the submitted work has not been published before (neither in English nor in any other language) and that the work is not under consideration for publication elsewhere.

## Acknowledgments

We thank the animal facility of the Max Planck Institute for Molecular Genetics, Berlin for expert support, especially Katja Zill and Ludger Hartmann. X.W. was supported by the Chinese Scholarship council (CSC), and the Sonnenfeld Stiftung Berlin. K.W and A.K. were supported by a grant from the German Ministry of Education and Research (BMBF), and the State of Brandenburg (DZD grant 82DZD00302).

## Online supplementary material

Table S1. Supporting Information (List of primers)

Data S1. Supporting Information (DESeq2 analysis of transcriptome data)

Data S2. Supporting Information (MaxQuant processed output files)

Data S3. Supporting Information (Differentially expressed proteins)

Figures S1 – S5. Supporting Information (Supplementary Figures)

