## Supplementary Figures and Tables for "Cell autonomous requirement of Neurofibromin (Nf1) for postnatal muscle hypertrophic growth and metabolic homeostasis"

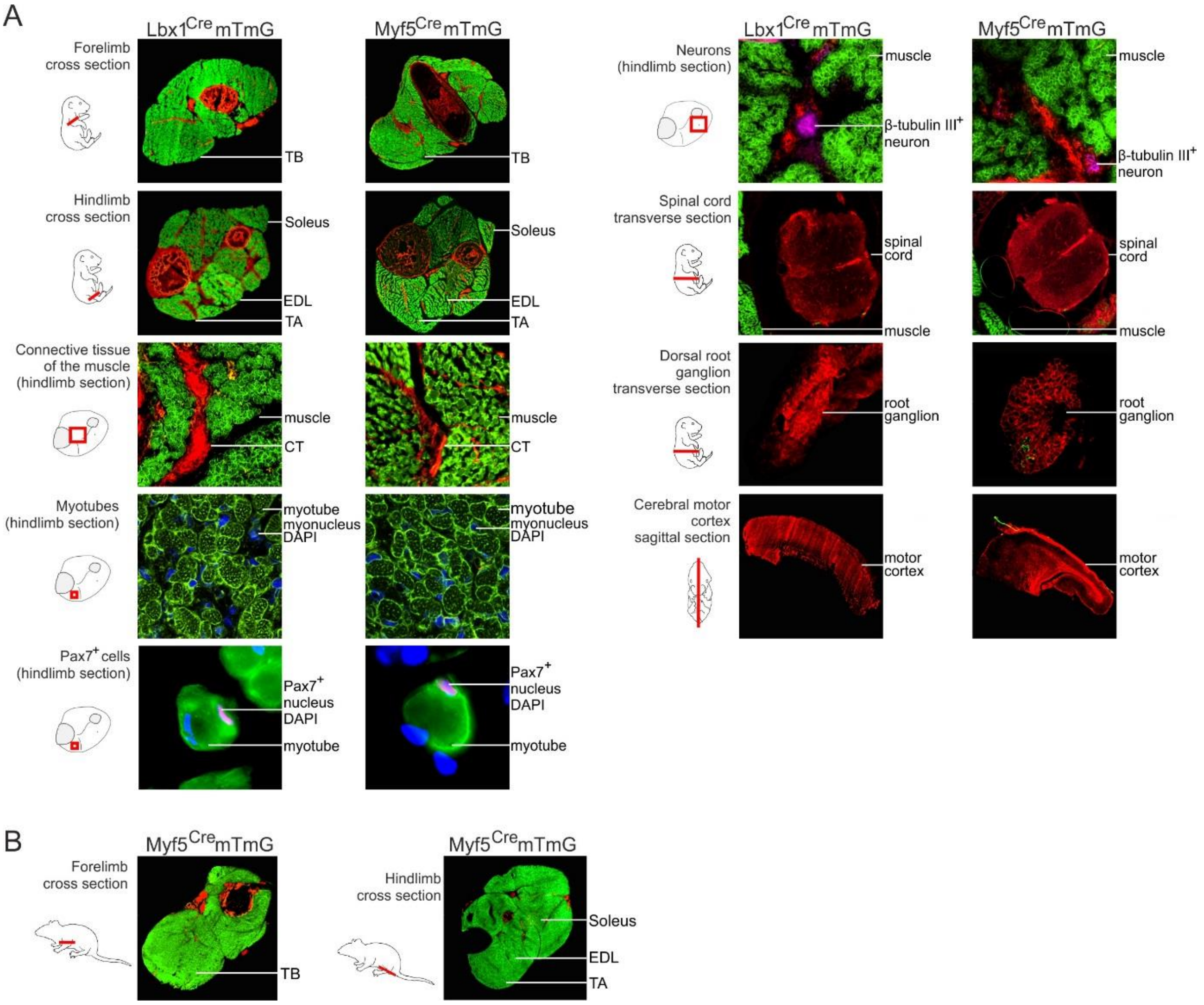


**Fig. S1** Characterization of Lbx1$^{Cre}$ and Myf5$^{Cre}$ specificity (A, B) Analysis of Lbx1Cre and Myf5Cre specificity. Cre mice were bred to Rosa26$^{mTmG}$ reporter mice and analyzed at embryonic day 18 (E18.5; A) or postnatal day 21 (p21; B). Staining: green depicts mG reporter activity tracing recombined cells, red depicts mT reporter activity tracing non-recombined cells. Satellite cells have been stained for Pax7, neurons for β-tubulin III (purple). TB: Triceps brachii; CT: connective tissue; EDL: Extensor digitorum longus; TA: Tibialis anterior.

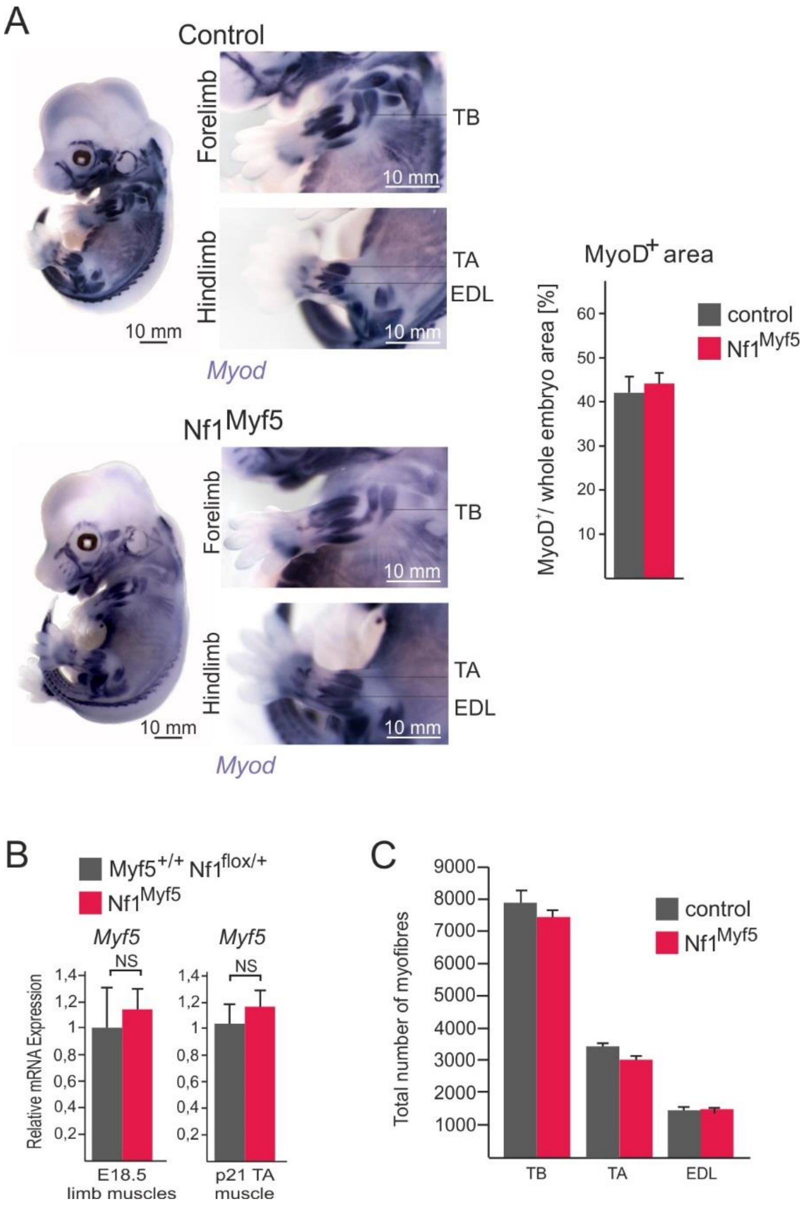


**Fig. S2** Extended data for pre- and postnatal characterization of Nf1$^{Myf5}$ animals (A) Whole-mount in-situ hybridization of 14 day old control and Nf1$^{Myf5}$ embryos for *Myod*. Quantification of *Myod*+ area in limbs is shown right (n=3 animals per genotype). (B) RT-qPCR analysis of *Myf5* expression from Myf5$^{+/+}$;Nf1$^{flox/+}$ vs. Myf5$^{Cre/+}$;Nf1$^{flox/flox}$ animals (n=3 animals per genotype). (C) Quantification of total myofiber numbers in muscles (TB: Triceps brachii, TA: Tibialis anterior, EDL: Extensor digitorum longus) of p21 control and Nf1$^{Myf5}$ animals (n=3 animals per genotype).

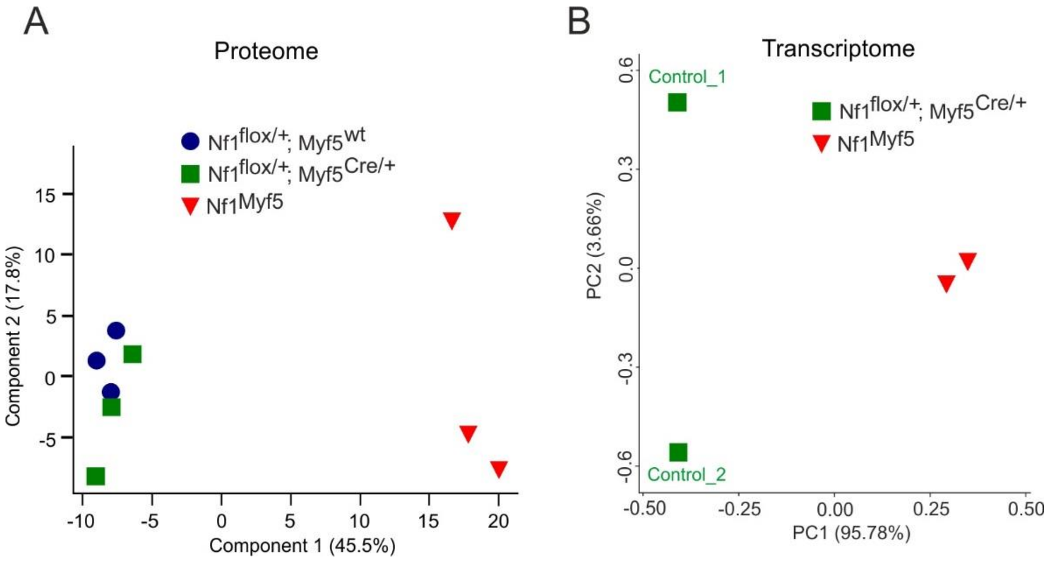


**Fig. S3** Extended data proteome and transcriptome analysis (A, B) Principal component analysis (PCA) of proteome (A) and transcriptome (B) data. Genotypes are indicated.

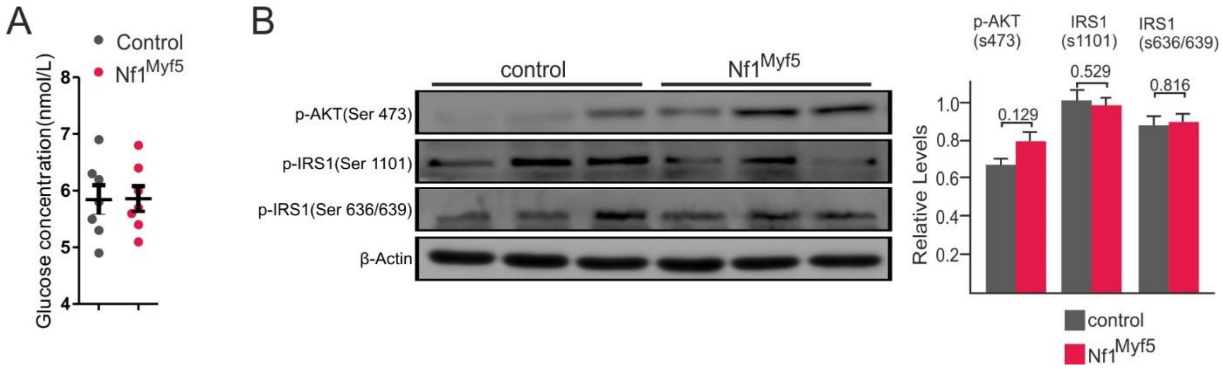


**Fig. S4** Blood glucose and insulin signaling in Nf1$^{Myf5}$ mice (A) Blood glucose test in 5 weeks old Nf1$^{Myf5}$ animals (n=7 animals per genotype; each dot represents measurement from one individual animal). (B) Western blot analysis for phosphorylated AKT (Ser473) and phosphorylated IRS1 (Ser1101, Ser 636/639). Quantification is shown right (n=3 animals per genotype).

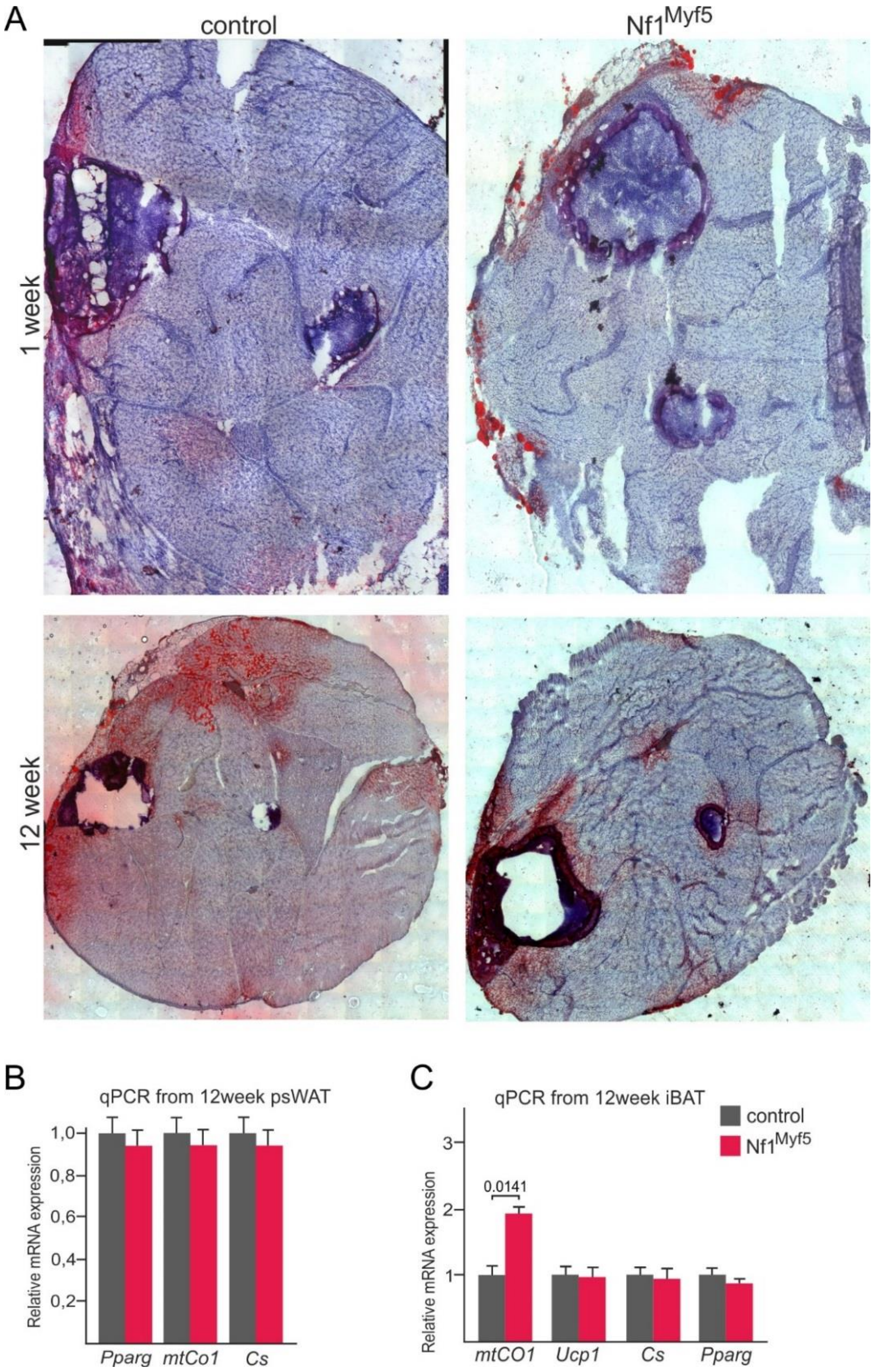


**Fig. S5** Extended data for analysis of adipose tissue in Nf1$^{Myf5}$ mice (A) Full images of hind limb cross sections stained for Oil red O and hematoxylin. Oil red O positive muscle-adjacent adipocytes can be seen, frequently lipids smear into the muscle area close to these sites. However the majority of muscle tissue is free of ectopic lipid staining in control and in Nf1$^{Myf5}$ mice. (B) RT-qPCR analysis of psWAT of of 12 week old control vs. Nf1$^{Myf5}$ mice for *Pparg*, *mtCO1* and *Cs* (n=3 animals per genotype). (C) RT-qPCR analysis of interscapular brown adipose tissue (iBAT) of 12 week old control and Nf1$^{Myf5}$ animals for *Ucp1*, *mtCO1*, *Cs* and *Pparg* mRNA (n=3 animals per genotype).

### Supporting Information Table S1

#### Genotyping primers

| *Primer Name* | *Primer Sequence* |
|---|---|
| *Lbx1_Cre_fw* | CGCCTTCCTCTCGCACCGTC |
| *Lbx1_Cre_rev* | GGCAGCCCGGACCGAC |
| *Myf5_Cre_fw* | CGTAGACGCCTGAAGAAGGTCAACCA |
| *Myf5_Cre_KO_rev* | ACGAAGTTATTAGGTCCCTCGAC |
| *Myf5_Cre_WT_rev* | CACATTAGAAAACCTGCCAACACC |
| *Nf1_P1* | CTTCAGACTGATTGTTGTACCTGA |
| *Nf1_P2* | CATCTGCTGCTCTTAGAGGAACA |
| *Nf1_P3* | ACCTCTCTAGCCTCAGGAATGA |
| *Nf1_P4* | TGATTCCCACTTTGTGGTTCTAAG |
| *Unspecific_Cre _fw* | GAGTGATGAGGTTCGCAAGA |
| *Unspecific_Cre_rev* | CTACACCAGAGACGGAAATC |

#### *Rt-PCR primers*

| *Primer Name* | Primer Sequence |
|---|---|
| *Myf5_fw* | TGAGGGAACAGGTGGAGAAC |
| *Myf5_rev* | CTGTTCTTTCGGGACCAGAC |
| *Nf1_fw* | ACAAAGGGTTACTGCCAT |
| *Nf1_rev* | GCTCCCCCAGATTTTTGC |
| *Myh2_fw* | ACCCTCCCAAGTACGACAAG |
| *Myh2_rev* | TACACCGGCAGCCATTTGTA |
| *Myh4_fw* | CAGAGTCACCTTCCAGCTCA |
| *Myh4_rev* | TGATTTCACCTTGACTGACGT |
| *Myh7_fw* | AGCTGGGAAGACTGTCAACA |
| *Myh7_rev* | CCAAAGGCCTCCAGAGCA |
| *Pparg_fw* | CGAGAAGGAGAAGCTGTTGG |
| *Pparg_rev* | TCAGCGGGAAGGACTTTATG |
| *Pgc1 α_fw* | AACCACACCCACAGGATCAGA |
| *Pgc1 α_rev* | TCTTCGCTTTATTGCTCCATGA |
| *Fabp3 _fw* | AGGTGGCTAGCATGACCAAG |
| *Fabp3_rev* | CTTGACCTTCCGGTCATCTG |
| *Fabp4_fw* | CTTTGCCACAAGGAAAGTGG |
| *Fabp4_rev* | GTCGTCTGCGGTGATTTCAT |
| *Acad1_fw* | GTAGCTTATGAATGTGTGCAACTC |
| *Acad1_rev* | GTCTTGCGATCAGCTCTTTCATTA |
| *Cpt2_fw* | GCTGCCTATCCCTAAACTTGAA |
| *Cpt2_rev* | CAATGCCGTTCTCAAAATCC |
| *Cpt1b_fw* | TCTGTGTCCGTCTCCTGTCC |
| *Cpt1b_rev* | CCATGCGGTAATATGCTTCA |
| *Idh3a_fw* | TCCTGGAGATGGAATTGGCC |
| *Idh3a_rev* | ATGGACTCCTTGGCTTCTGG |
| *Lpl_fw* | TCATCTCATTCCTGGATTAGCA |
| *Lpl_rev* | GGCCCGATACAACCAGTCTA |
| *Il6_fw* | TGAACAACGATGATGCACTTG |
| *Il6_rev* | TCTCTCTGAAGGACTCTGGC |
| *Fgf21_fw* | ACCTGGAGATCAGGGAGGAT |
| *Fgf21_rev* | GAGAGCTCCATCTGGCTGTT |
| *Fgf23_fw* | TACAGCCAGGACCAGCTATC |
| *Fgf23_rev* | TGGCTCCTGTTATCACCACA |
| *Fgf15_fw* | CGCTACTCGGAGGAAGACTG |
| *Fgf15_rev* | TTGGCCTGGATGAAGATGAT |
| *Adipoq_fw* | GGAGATGTTGGAATGACAGGA |
| *Adipoq_rev* | CGAATGGGTACATTGGGAAC |

| | |
|---|---|
| *Gapdh-fw* | AACTTTGGCATTGTGGAAGG |
| *Gapdh-rev* | CAGTCTTCTGGGTGGCAGTG |
| *Actb-fw* | CGTGAAAAGATGACCCAGATCA |
| *Actb-rev* | GGGACAGCACAGCCTGGAT |
| *4ebp1-fw* | GGGGACTACAGCACCACTCC |
| *4ebp1-rev* | ATCGCTGGTAGGGCTAGTGA |
| *Ctsl-fw* | TTAGTGCAGAGTGGCACCAG |
| *Ctsl-rev* | CCGTTGTGTAGCTGGATCATT |
| *Psma-fw* | GCCGCTCACCAGAAGAAAAT |
| *Psma-rev* | GACGAGACACAGGAAGTGGTC |
| *Fbxo32-fw* | CGCCATGGATACTGTACTTTG |
| *Fbxo32-rev* | TGAAGTTCTTTTGGGCGATG |
| *Trim63-fw* | ACTGCATCTCCATGCTGGT |
| *Trim63-rev* | CAGCTCGCTCTTCTTCTCGT |
| *Gabarapl1-fw* | TCGTGGAGAAGGCTCCTAAA |
| *Gabarapl1-rev* | GTCCTCAGGTCTCAGGTGGA |
| *Nfe2l2-fw* | CCCACATTCCCAAACAAGAT |
| *Nfe2l2-rev* | GAGGGGCAGTGAAGACTGAA |
| *Ucp1-fw* | GTCAGAATGCAAGCCCAGAG |
| *Ucp1-rev* | ACCAGCTCTGTACAATTGATGA |
| *Sod1-fw* | AAGCGGTGAACCAGTTGTGT |
| *Sod1-rev* | GGGCCACCATGTTTCTTAGAG |
| *Cs-fw* | CCCAGGATACGGTCATGCA |
| *Cs-rev* | GCAAACTCTCGCTGACAGGAA |
| *mtCO1-fw* | ACATGAAACCCCCAGCCATA |
| *mtCO1 -rev* | TCCAAATCCTGGGAGGATAA |
